# Polyamine-mediated sensitization of *Klebsiella pneumoniae* to macrolides through a dual mode of action

**DOI:** 10.1101/2024.02.18.580908

**Authors:** Joshua M. E. Adams, Peri B. Moulding, Omar M. El-Halfawy

## Abstract

Chemicals bacteria encounter at the infection site could shape their stress and antibiotic responses; such effects are typically undetected in standard lab conditions. Polyamines are small molecules typically overproduced by the host during infection and have been shown to alter bacterial stress responses. We sought to determine the effect of polyamines on the antibiotic response of *Klebsiella pneumoniae*, a Gram-negative priority pathogen. Interestingly, putrescine and other natural polyamines sensitized *K. pneumoniae* to azithromycin, a macrolide protein translation inhibitor typically used for Gram-positive bacteria. This synergy was further potentiated in the physiological buffer, bicarbonate. Chemical genomic screens suggested a dual mechanism whereby putrescine acts at the membrane and ribosome levels. Putrescine permeabilized the outer membrane of *K. pneumoniae* (NPN and β-lactamase assays) and the inner membrane (*Escherichia coli* β-galactosidase assays). Chemically and genetically perturbing membranes led to a loss of putrescine-azithromycin synergy. Putrescine also inhibited protein synthesis in an *E. coli*-derived cell-free protein expression assay simultaneously monitoring transcription and translation. Profiling the putrescine-azithromycin synergy against a combinatorial array of antibiotics targeting various ribosomal sites suggested that putrescine acts as tetracyclines targeting the 30S ribosomal acceptor site. Next, exploiting the natural polyamine-azithromycin synergy, we screened a polyamine analog library for azithromycin adjuvants, discovering four azithromycin synergists with activity starting from the low micromolar range and mechanisms similar to putrescine. This work sheds light on the bacterial antibiotic responses under conditions more reflective of those at the infection site and provides a new strategy to extend the macrolide spectrum to drug-resistant *K. pneumoniae*.

---

Antimicrobial resistance (AMR) is an increasingly rising problem gravely impacting today’s society. In 2019, there was an estimated 4.95 million deaths worldwide associated with bacterial AMR ^1^. The AMR public health crisis prompted the World Health Organization (WHO) to issue a list of priority pathogens posing the most serious threat that requires urgent action; Gram-negative bacteria make up the majority of this list due to the limited therapeutic options available against these bacteria^2–4^. Unlike Gram-positive bacteria, the Gram-negative ones possess an outer membrane, an integral component of which are lipopolysaccharides (LPS), which constitutes a formidable barrier that prevents antibiotic permeation ^5^. *Klebsiella pneumoniae* is one of the Gram-negative bacteria on the WHO priority pathogens list, gaining international concern due to the high prevalence of hypervirulent and drug resistant strains ^4^. With the constant rise in AMR and the inadequate antibiotic drug discovery pipeline ^6^, there is a dire need for new strategies to combat Gram-negative resistant strains^1^.

Bacteria encounter a myriad of chemicals at the infection site that may alter their antibiotic susceptibility and virulence ^7, 8^. The potential effects of such chemicals are not typically captured in standard antibiotic susceptibility assays, whose conditions follow the guidelines issued by standardizing agencies such as the Clinical and Laboratory Standards institute (CLSI). Polyamines constitute an important class of chemicals present at the infection sites, especially since they are overproduced by the host during infection ^9^. Natural polyamines, such as spermine, spermidine, and putrescine, are small polycationic molecules produced by almost all living organisms and are critical for their viability ^10^. During infection, polyamine biosynthesis is upregulated in regenerating tissues of the host at the inflammatory and infection site; for example, polyamines accumulate in the lungs during pneumonia infections, suggesting a role in suppressing infection ^11–13^. Additionally, polyamines play an immunomodulatory role, where spermine exhibits macrophage suppression and anti-inflammatory activity ^12^. The polyamine backbone structure consists of regularly spaced positive charges interrupted by hydrophobic hydrocarbon bridges; this unique structure allows polyamines to interact with various macromolecules ^14^. Polyamines can stabilize nucleic acids and thus influence transcription, chromosomal structure, apoptosis, and protein synthesis ^15–20^; they are involved in cell growth, differentiation, cell membrane stability and immune responses in humans ^14, 15^.

Polyamines are similarly involved in cellular homeostasis and stress response in bacteria^21^. They can also alter antibiotic susceptibility in a wide range of Gram-positive and Gram-negative bacteria ^22–27^. For example, polyamines increased resistance of *Pseudomonas aeruginosa* and *Burkholderia cenocepacia* to antibiotics, including polymyxins and fluoroquinolones ^23–25^, and conversely increased the susceptibility of *Staphylococcus aureus* and enteric bacteria, such as *Escherichia coli*, to β-lactam antibiotics ^26^.

Macrolides are a class of antibiotics that target and inhibit protein synthesis via binding to the 23S portion of the 50S bacterial ribosome subunits^28^. Antibiotics in this class are highly prescribed due to their potency, low toxicity, and favourable pharmacokinetic properties, particularly their oral bioavailability ^29–31^. Newer classes of macrolides, such as the azalide azithromycin, are clinically advantageous due to their extensive and rapid distribution into tissues and intracellular compartments yet minimal accumulation in fat and muscle ^30^. Typically, macrolides are used to treat Gram-positive infections with less application in Gram-negative ones due to their poor penetration of the outer membrane ^32^. However, macrolides show potency against Gram-negative bacteria albeit under certain infection-relevant conditions. For example, *E. coli* exhibit heightened sensitivity to macrolides in the presence of the physiologically relevant buffer bicarbonate, which interferes with the proton motive force increasing transport of macrolides through the inner membrane ^33^. Additionally, azithromycin treatment has been shown to improve the therapeutic outcome of cystic fibroses (CF) patients infected with the Gram-negative pathogen *P. aeruginosa* ^34^, albeit with an unknown mechanism. Interestingly CF patients with pulmonary exacerbations can exhibit an 8-fold increase in polyamine levels ^35^. Understanding the effects of macrolides against priority Gram-negative bacteria, such as *K. pneumoniae*, under host relevant conditions, especially in the presence of polyamines, may expand their spectrum of activity to this group of bacteria where the clinical need is high.

In this study, we show that polyamines sensitize *K. pneumoniae* to the macrolide azithromycin. We determined the mechanism of this synergy, revealing both a membrane and target-dependent effects. These observations prompted us to screen a collection of synthetic polyamine analogs for macrolide adjuvants against *K. pneumoniae*, identifying micromolar azithromycin synergists. Together, this work offers potential new antimicrobial strategies against multi-drug resistant Gram-negative bacteria.

## Results and discussion

### Natural polyamines synergize with azithromycin

We conducted checkerboard assays to test the effect of the polyamine putrescine on azithromycin susceptibility in *K. pneumoniae* MKP103, a derivative of the clinical outbreak isolate KPNIH1 lacking its carbapenemase-encoding gene ^36^. Putrescine synergized with azithromycin with a fractional inhibitory concentration index (FICI) of 0.47±0.01 (Fig 1A). FICI ≤ 0.5 were interpreted as synergy, FICI > 4.0 as antagonism, and 0.5 < FICI ≤4.0 as additive, indifference, or no interaction. This synergy matches previously reported polyamine-induced macrolide sensitivity in another Gram-negative organism, *E. coli* ^26^; the mechanism of such interaction is unknown. The putrescine-azithromycin synergy reversed macrolide resistance of *K. pneumoniae* MKP103; putrescine at 20 mM lowered the azithromycin MIC to below the CLSI clinical breakpoint (Fig 1A). A bacterial strain is deemed sensitive to an antibiotic at MIC values less than the clinical breakpoint, thus may become clinically used^37^. We also observed azithromycin synergy with other natural polyamines (spermine and spermidine, Fig 1B-C). Polyamine physiological levels vary but are generally within the millimolar range and are further increased at the infection site upon infection ^38, 39^; hence polyamines at the site of infection may be sufficient to drive azithromycin synergy similar to what we observed *in vitro*.

**Figure 1.**
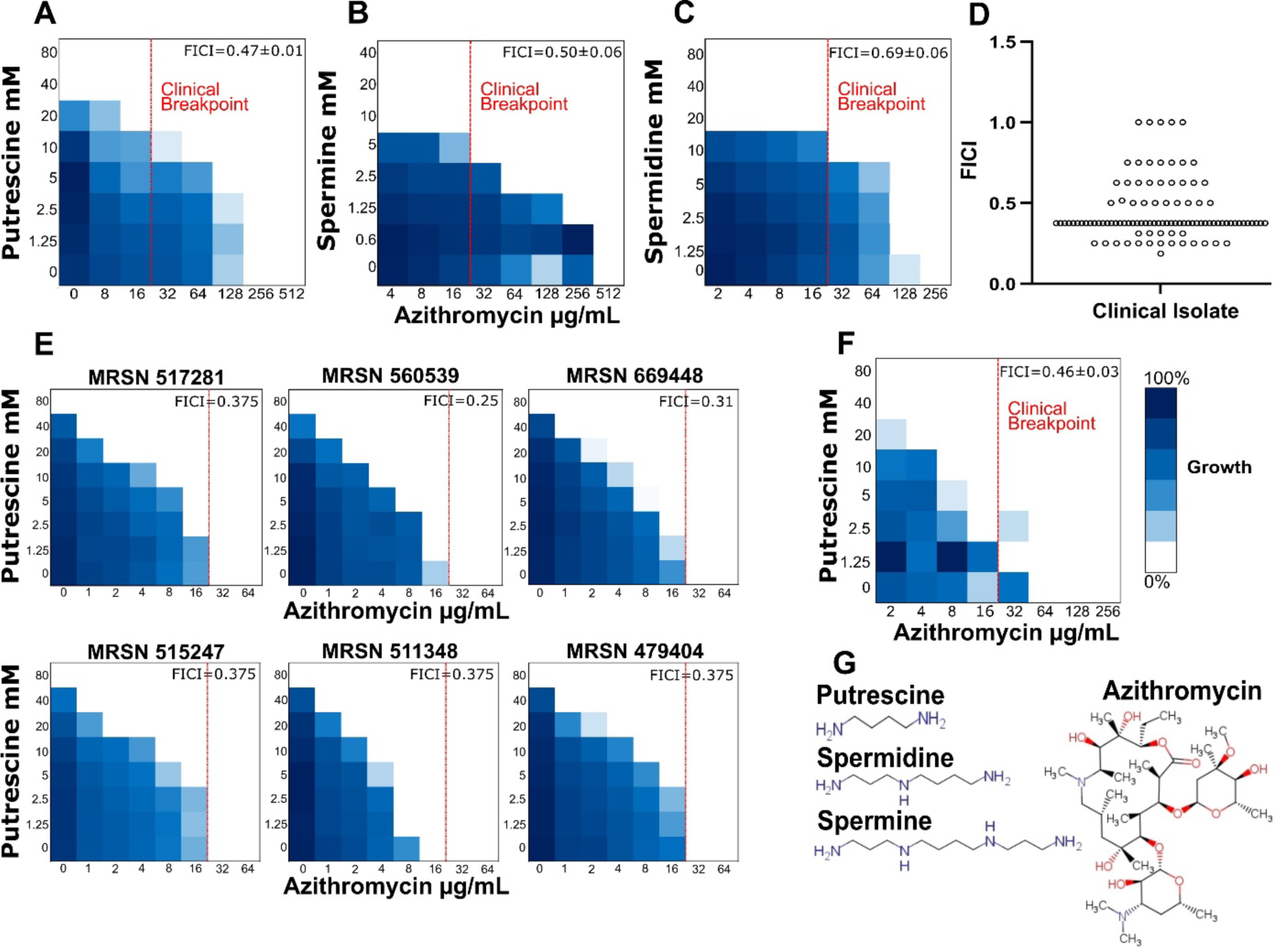
Natural polyamines synergize with azithromycin against *K. pneumoniae* rendering it susceptible to the macrolide antibiotic. A-C) Representative checkerboard assays of *K. pneumoniae* MKP103 of azithromycin combined with putrescine, spermine, and spermidine. D) A scatter plot representing the distribution of FICI values across the *K. pneumoniae* diversity panel. E) Checkerboard assays of select representative clinical isolates from the diversity panel (n=1 for each isolate, results of the rest of the 100 isolates are shown in Table S1). F) A checkerboard assay of putrescine and azithromycin against *K. pneumoniae* MKP103 in the presence of 25 mM bicarbonate buffer. G) Chemical structures of the polyamines used in this study and azithromycin. Dark blue regions represent high cell density determined by OD_600._ Red dotted lines refer to the clinical breakpoint as set by CLSI. Mean of the FICI and SEM (standard error of the mean) are displayed on each representative checkerboard, n≥3.

The polyamine-azithromycin synergy was maintained across diverse *K. pneumoniae* clinical isolates. We conducted putrescine-azithromycin checkerboard assays against the *K. pneumoniae* diversity panel, comprising 100 clinical isolates collected from 63 facilities in 19 countries with an array of genotypic and phenotypic differences that span pan-sensitive, pan-resistant, and hypervirulent lineages (Table S1) ^40^. We observed the putrescine-azithromycin synergy in 78 isolates (Fig 1D), which lowered azithromycin MIC below the CLSI breakpoint in 77 isolates (Table S1; examples in Fig 1E). Notably, such synergy was conserved across pan-sensitive to pan-drug-resistant isolates, including those encoding β-lactamases (including CTX-M β-lactamases and NDM and OXA carbapenemases), Rmt(B,F,H) methyltransferases, MCR-1, Tet(X), and ErmB, which drive resistance against β-lactams (including carbapenems), aminoglycosides, colistin, tetracyclines, and erythromycin, respectively (Table S1). On the other hand, the putrescine-azithromycin combination showed additive effects in 22 isolates, which harbour genes encoding the macrolide phosphotransferases Mph(A), Mph(B), and Mph(E) (Table S1).

Next, we sought to test whether polyamines further sensitize *K. pneumoniae* to azithromycin in the presence of bicarbonate, the physiological buffer, in which Gram-negative bacteria are less resistant to macrolides ^33^. We show that exposure to the physiological bicarbonate concentration (25 mM) resulted in 4-fold reduction in azithromycin MIC against *K. pneumoniae* (Fig 1F compared to 1A), as previously reported ^33^. Putrescine further sensitized *K. pneumoniae* to azithromycin (further 16-fold reduction in MIC) in the presence of bicarbonate (Fig 1F). These observations suggest that multiple physiologically-relevant factors may act in tandem to alter bacterial responses to antibiotics.

### Genomic screens identify a dual mechanism of the polyamine-azithromycin synergy

To identify the mechanism of polyamine-azithromycin synergy, we hypothesized that a mutant in the gene encoding the determinant of synergy will show increased azithromycin susceptibility and loss in synergy with polyamines. We used an arrayed sequence-defined transposon mutant library covering the non-essential genome of *K. pneumoniae* MKP103, comprising one or multiple non-redundant mutants for each non-essential gene, with an average of 2.5 mutants per gene ^36^. First, we screened the library at two azithromycin sub-inhibitory concentrations, 32 and 64 µg/mL, to identify mutants susceptible to azithromycin (Fig 2A-B and Fig S1). The screen revealed 71 mutants representing 67 genes showing partial or complete growth inhibition at the sub-inhibitory azithromycin concentrations (Fig 2A-B and Fig S1).

**Figure 2.**
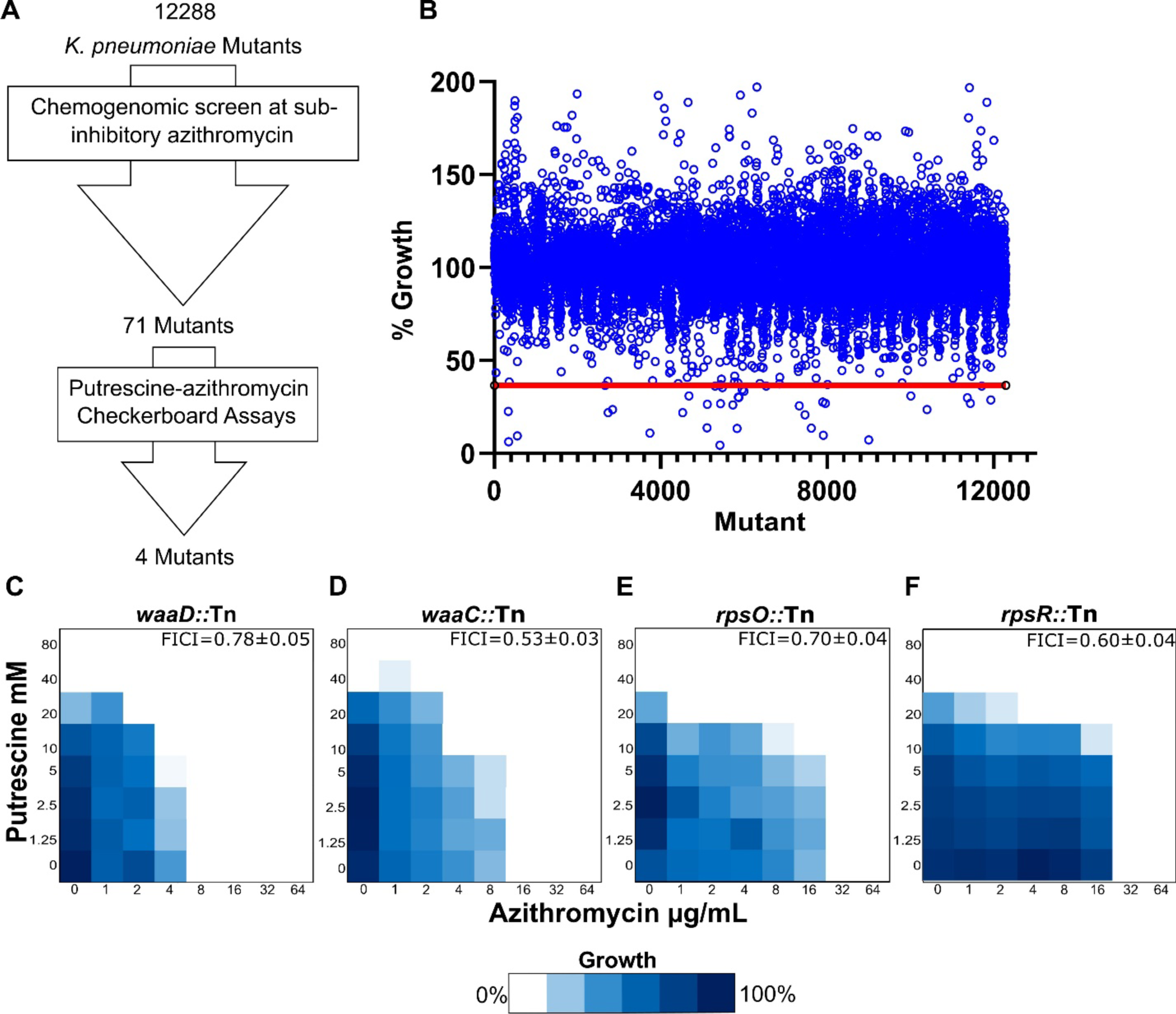
A chemogenomic screen identifies putrescine-azithromycin synergy determinants. A) A flow chart of the chemogenomic screen process that led to the identification of genetic determinants of azithromycin-putrescine synergy B) An index plot of the primary screen of *K. pneumoniae* MKP103 mutants grown in 32 μg/mL of azithromycin shown as % growth relative to that of the untreated mutants. Data points below a cut-off (the red line) of 1.5x the standard deviation of all datapoints below the interquartile mean are considered putative hits. C-F) Heat maps of representative putrescine-azithromycin checkerboards of the *K. pneumoniae* MKP103 mutants identified from the chemogenomic screen. Dark blue represents maximal growth determined by OD_600_. Mean of the FICI and SEM are displayed on each representative checkerboard, n=3.

These genes include those whose products are involved in stress induced mutagenesis (*recC* and *recB*), ribosomal function (*rpsO, rpsR, ybeY,* and *rsmB*), and membrane integrity (*waaD, waaC, waaE, dsbA, acrB,* and *bamB*); knockouts of genes involved in these functions have been previously identified in either *K. pneumoniae* or *E. coli* or both as more susceptible to macrolides and other protein translation inhibitors ^41–43^.

Next, we conducted putrescine-azithromycin checkerboard assays against the 71 mutants (Fig S2 and Fig 2C-F). Azithromycin sensitivity (MIC ≤ 32µg/mL) and lack of synergy (FICI >0.5), suggested mutants of genetic determinants of the putrescine-azithromycin synergy; mutants with disruption in four genes (*waaD, waaC*, *rpsO,* and *rpsR*) fit these criteria (Fig 2C-F). Of note, none of the mutants exhibited a completely additive phenotype showing moderate potentiation below the cut-off for synergy, suggesting a potential involvement of multiple determinants. The disruptions in these mutants can be categorized into membrane (*waaD*::Tn and *waaC*::Tn) and 30S ribosomal subunit (*rpsO*::Tn and *rpsR*::Tn) related disruptions.

Knockouts of *waaD* and *waaC* lead to the same deep rough LPS phenotype ^44–46^. WaaD is necessary for the production of ADP-L-glycero-D-mannoheptose, which WaaC incorporates into the LPS structure; loss of these enzymes results in a truncated LPS without an outer core and most of the inner core exposing Kdo2-lipid A (Fig S3A-B)^44^. Notably, the *waaD*::Tn was hypersensitive to novobiocin (Fig S4), as expected for LPS deep rough mutants ^44, 47^. The azithromycin hypersensitivity of these *K. pneumoniae* mutants is in agreement with previous reports of *E. coli* with a compromised LPS structure being more sensitive to macrolides ^41, 48^.

Setting the azithromycin susceptibility cutoff to MIC ≤64µg/mL led to the inclusion of *dsbA*::Tn, *bamB*::Tn, and KPNIH1_25560::Tn (predicted *waaE* ^46^), which exhibited partial loss of synergy (Fig S2A-C and Fig S3C). Loss of DsbA leads to accumulation of inactive LptD (a lipopolysaccharide assembly protein), BamB is a lipoprotein essential for assembly and insertion of β-barrel proteins into the outer membrane, and WaaE is responsible for the transfer of the branch β-D-glucopyranose (Glc) to L-glycero-D-manno-heptose I (Heptose I) ^46, 49, 50^. Loss of *dsbA* and *waaE* has been previously shown to exhibit macrolide sensitivity in *E. coli* and *K. pneumoniae,* respectively ^43, 46^, whereas knockouts of *bamB* exhibit loss of a membrane barrier effect in *E. coli,* becoming sensitive to novobiocin ^50^.

We tested other mutants involved in LPS biosynthesis to determine the extent of LPS truncation necessary for the loss of putrescine-azithromycin synergy. WaaF and WaaQ are enzymes responsible for the sequential addition of heptose II and heptose III, respectively, to complete the LPS inner core ^46^. Knockouts of genes encoding both enzymes showed no change in azithromycin MIC and retained putrescine-azithromycin synergy (Fig S3D-E). Together, a deep rough truncation of the LPS core leads to loss of putrescine-azithromycin synergy, suggesting that a similar polyamine-mediated membrane permeabilization induces the observed synergy (Fig S3). For subsequent follow up assays, we used *waaD*::Tn as a representative of the deep rough mutants.

The ribosomal proteins RpsO, RpsR, and RpsF create a complex within the platform domain of the 30S ribosomal subunit, where RpsR and RpsF form heterodimers ^51^. Both *rpsO*::Tn and *rpsR*::Tn mutants exhibited an azithromycin MIC of 32 µg/mL and an FICI value of 0.7 ± 0.04 and 0.6 ± 0.04, respectively (Fig 2G-H). RpsF is not represented in the *K. pneumoniae* MKP103 library. The deletion of *rpsO* has been reported to influence the functional capacity of 30S subunits and decrease the amount of 70S complexes ^52^. RpsR is hypothesized to be involved in the binding of aminoacyl-tRNA to the P site of the ribosome ^53^. Genes encoding other 30S ribosomal subunit proteins are essential except for *rpsA,* whose mutant yielded no change in azithromycin susceptibility (data not shown). RpsA is weakly associated with the 30S ribosome and even lacks association to the ribosome in some *E. coli* strains ^54^. Notably, the checkerboard of *infB*::Tn, a mutant of the translation initiation factor for *rpsO*, exhibited a partial reduction in synergy and a 4-fold decrease in azithromycin MIC compared to that of the wild type (Fig S2D).

Together, our results suggest that polyamines synergize with azithromycin via a dual mechanism of action: (1) increasing membrane permeability allowing azithromycin to cross this barrier, and (2) acting on the 30S ribosomal subunit compounding the effects of azithromycin in suppressing protein synthesis.

### Putrescine permeabilizes bacterial outer and inner membranes

Gram-negative bacteria, including *K. pneumoniae*, possess outer and inner membranes. We studied the effects of polyamines on both membranes. Starting with the outer membrane, we showed, in kinetic 1-*N*-phenylnaphthylamine (NPN) fluorometric assays, that putrescine disrupted the outer membrane of *K. pneumoniae* shown as an increase in NPN uptake (Fig 3A). A disruption of the outer membrane occurs at 10 mM and 20 mM of putrescine (P < 0.0001), which is within the synergistic range of the putrescine-azithromycin interaction. Such putrescine-mediated outer membrane perturbation was similar to that displayed by the deep rough LPS mutant *waaD*::Tn, which showed a similar increase in NPN uptake relative to the wild type (P < 0.005, Fig 3B). *Salmonella enterica* serovar Typhimurium and *E. coli waaD* deletion mutants also showed a similar (2 to 5-fold) increase in NPN uptake in comparison to the respective parent strains ^55, 56^. Using colorimetric quantification of the hydrolysis of nitrocefin, a β-lactam probe, by a periplasmic β-lactamase constitutively expressed from pBR322 as another proxy for outer membrane perturbation, we further confirmed the increased outer membrane permeability of the *waaD*::Tn mutant in comparison to its parent strain (Fig 3C), in agreement with previous reports^57^. Given that outer membrane permeabilization of the deep rough mutant (Fig 3B-C) is sufficient to enable macrolides to cross that barrier (Fig 1A, Fig 2C-D, ^41^), further membrane perturbation may not lead to enhancement of the macrolide activity. Thus, the lack of putrescine-azithromycin synergy in *waaD*::Tn further suggests that the polyamine-mediated outer membrane perturbation contributes to the synergy. As such, we reasoned that chemically or genetically perturbing the membrane should exhibit increased susceptibility to azithromycin and a loss of synergy with putrescine, similar to *waaD*::Tn. Chemically perturbing *K. pneumoniae* membrane by 1 µg/mL of polymyxin B nonapeptide (PMBN), a cationic peptide that disrupts the outer membrane with little to no antimicrobial activity ^58^, led to a 4-fold decrease in azithromycin MIC (Fig 3 D-E), consistent with previous reports ^5, 58, 59^, and a loss of putrescine-azithromycin synergy (FICI 0.83 ± 0.10) (Fig 3E). A moderate potentiation of azithromycin (4-fold MIC shift, Fig 3E) in the presence of 20 mM putrescine (below the cut-off for synergy) may be the result of the dual mechanism of action of putrescine. Similarly, an *E. coli* with a compromised outer membrane genetically-induced by the deletion of 9 efflux pump transmembrane proteins (Δ9 mutant) ^60^, exhibited an 8-fold decrease in azithromycin MIC and a loss in synergy (FICI = 1.00 ± 0.00) relative to its parent strain (M6394), which showed a synergistic phenotype (FICI = 0.41 ± 0.07) (Fig S5A-B). The *E. coli* wild type and Δ9 mutant showed NPN profiles (Fig S5C) similar to those of *K. pneumoniae* MKP103 wild type and *waaD*::Tn (Fig 3B).

**Figure 3.**
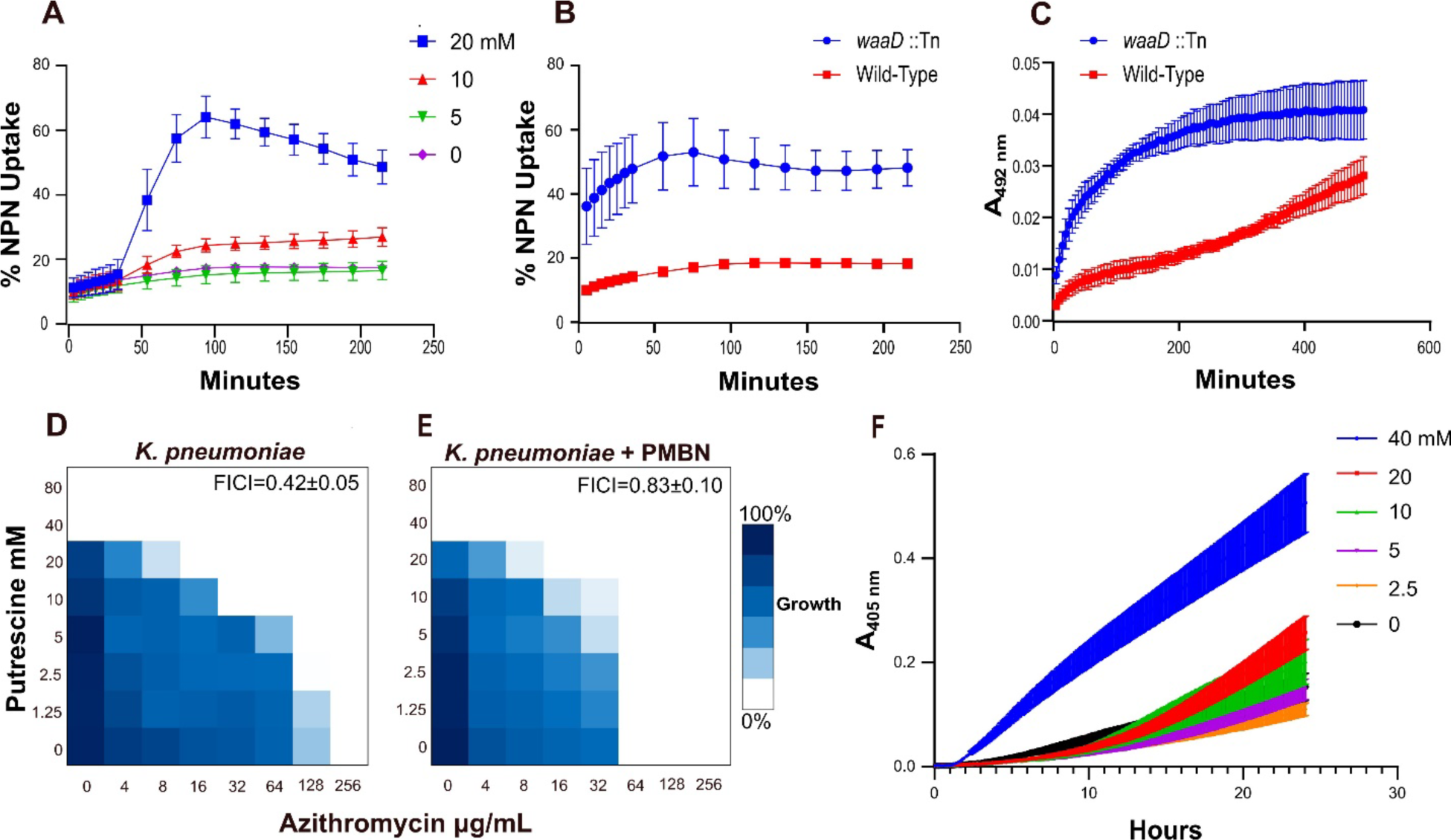
Putrescine permeabilizes bacterial outer and inner membranes. A-B) Fluorometric kinetic assays displaying % NPN uptake in wild type *K. pneumoniae* MKP103 treated with different concentrations of putrescine (A) and in *K. pneumoniae waaD*::Tn relative to its parent MKP103 strain (B). C) A kinetic absorbance assay displaying nitrocefin hydrolysis by a periplasmic β-lactamase in *K. pneumoniae* wild type and *K. pneumoniae waaD*::Tn. D-E) Representative checkerboards of putrescine-azithromycin combination against *K. pneumoniae* MKP103 in the absence (D) and presence (E) of 1 µg/mL polymyxin B nonapeptide; dark blue regions represent high cell density determined by OD_600_. F) A kinetic absorbance assay of ONPG hydrolysis in *E. coli* ML-35 as a proxy for inner membrane perturbation. All kinetic assays are the combined data of three independent assays each done in triplicate. All error bars denote SEM. Mean of the FICI and SEM are displayed on each representative checkerboard, n=3.

Notably, spermidine and spermine at the micromolar range have been reported to confer resistance to polymyxin B in *P. aeruginosa* by localizing at the outer membrane. Molecular dynamics simulations of model bacterial outer membranes have shown that polyamines engage in electrostatic interactions with negatively charged phosphate groups, stabilizing the outer membrane with the largest stabilizing effects in the order of spermidine, spermine, then putrescine ^61^. The activity of putrescine at the outer membrane in *K. pneumoniae* at the physiologically relevant millimolar range has not been previously reported. At low putrescine concentrations, we confirmed that putrescine can stabilize the outer membrane shown as reduced NPN uptake relative to the untreated cells (Fig S6). As such, it appears that putrescine displays a dose-dependent activity on the outer membrane, ranging from stabilization at low concentrations to disruption at higher concentrations.

The inner membrane of Gram-negative bacteria may constitute a second physical barrier for azithromycin. We sought to test if putrescine also affects the inner membrane by measuring the β-galactosidase-mediated o-nitrophenyl-β-D-galactopyranoside (ONPG) hydrolysis in a lactose permease-deficient *E. coli* that constitutively expresses a cytoplasmic β-galactosidase (*E. coli* ML-35). Putrescine at 40 mM significantly increased ONPG hydrolysis (P < 0.0001, Fig 3F), suggesting that polyamines can also disrupt the inner membrane, potentially contributing to increasing intracellular azithromycin concentration. It has been previously shown that disrupting the inner membrane in Gram-negative bacteria lowers the MIC of azithromycin ^33^. Taken together, we have shown evidence that putrescine permeabilizes both the outer and inner membranes, contributing to its synergy with azithromycin.

### Polyamines act on the ribosome compounding the protein synthesis inhibitory effect of azithromycin

The lack of azithromycin-putrescine synergy in the 30S ribosomal subunit mutants (*rpsO*::Tn and *rpsR*::Tn, Fig 2E-F) suggests the synergy may be driven, at least in part, by an interaction between both molecules at the ribosomal level. Hence, we sought to test the effect of putrescine on protein translation. To this end, we used a cell-free protein synthesis system, an extract-based transcription/translation system derived from *E. coli* designed to synthesize proteins under the control of T7 RNA Polymerase. We quantified the levels of a plasmid-encoded destabilized enhanced Green Fluorescent Protein (deGFP) fluorometrically in the presence and absence of putrescine (Fig 4 and Fig S7). Simultaneously, we measured the effects on transcriptional activity in real-time (Fig 4 and Fig S7A-C), since the plasmid also included a malachite green mRNA aptamer allowing for such measurement fluorometrically via binding of malachite green dye ^62^. At the higher tested concentration of putrescine, mRNA levels were reduced but protein synthesis was further inhibited (Fig S7A-F). Similar reduction in gene expression has been previously observed in cell-free *in vitro* gene expression models at high spermine and spermidine concentrations ^63^. Thus, to determine the effect of putrescine on protein synthesis due to a ribosomal effect, the deGFP signal was normalized by the mRNA signal (Fig 4). The lower putrescine concentrations exhibited increased protein synthesis (in the first ∼90 minutes of exposure, Fig 4), which is in agreement with previous reports that low concentrations of polyamines promote protein synthesis in cell free systems ^63^. A majority of intracellular polyamines exist as polyamine-RNA and polyamine-ribosome complexes with evidence suggesting they contribute to transcription and translation, playing an essential role in ribosome biogenesis, tRNA positioning on the ribosome, and translation elongation speed ^64–67^. Cross-linking studies showed that ribosomes have >500 stably linked polyamines per ribosome, and it has been suggested polyamine binding happens in a stochastic manner ^67, 68^. On the other hand, our results show that 80 mM of putrescine marked a significant decrease in protein synthesis, suggesting a decreased ribosomal function of the cell-free protein expression system (P=0.0001, Fig 4). The addition of spermine to ribosome complexes was previously shown to slow down protein expression, inhibit peptidyl transferase activity, and possibly alter ribosome conformation^69, 70^.

**Figure 4.**
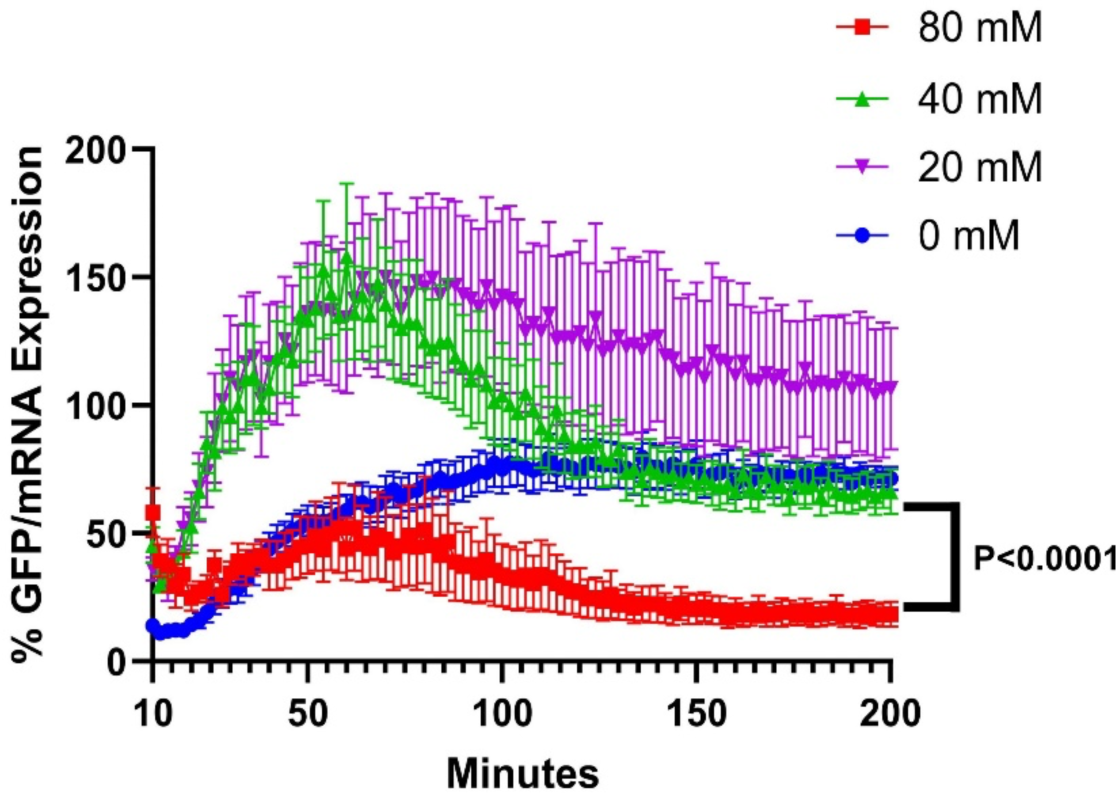
Putrescine inhibits protein synthesis. Protein (deGFP) and mRNA levels were simultaneously monitored fluorometrically (433/475 and 610/650 nm, respectively) in black 384 well plates using the NEBExpress^TM^ cell-free protein expression system in 200 mM HEPES buffer with an engineered plasmid encoding deGFP with a malachite green mRNA aptamer. Results shown are normalized by dividing each well’s RFU deGFP value by the malachite green RFU value and converted into a percentage of the maximal signal observed in the untreated condition. Raw RFU data found in Fig S7A-F. All error bars denote SEM. A one-way ANOVA analysis followed by a Brown-Forsythe and Welch test assessed the significance of the difference as indicated on the plot.

Next, we sought to determine the potential target site of putrescine within the ribosome through a phenotypic combinatorial approach using antibiotics targeting different ribosomal sites and a control (rifampicin) acting outside the ribosome. We hypothesize that, in the wild type, antibiotics targeting the same site within the ribosome as that of putrescine will show similar macrolide synergy as that of putrescine-azithromycin combination. Additionally, we hypothesize that a compound that mirrors the phenotype of putrescine-azithromycin will show additive effects when in combination with putrescine. We tested eight antibiotics, including azithromycin, in combination with one another and with putrescine to compare their FICI values (Fig 5A). These antibiotics targeted various sites within the ribosome: Paromomycin and gentamicin bind to the aminoacyl site within the 30S ribosomal subunit leading to misreading of genetic code.

**Figure 5.**
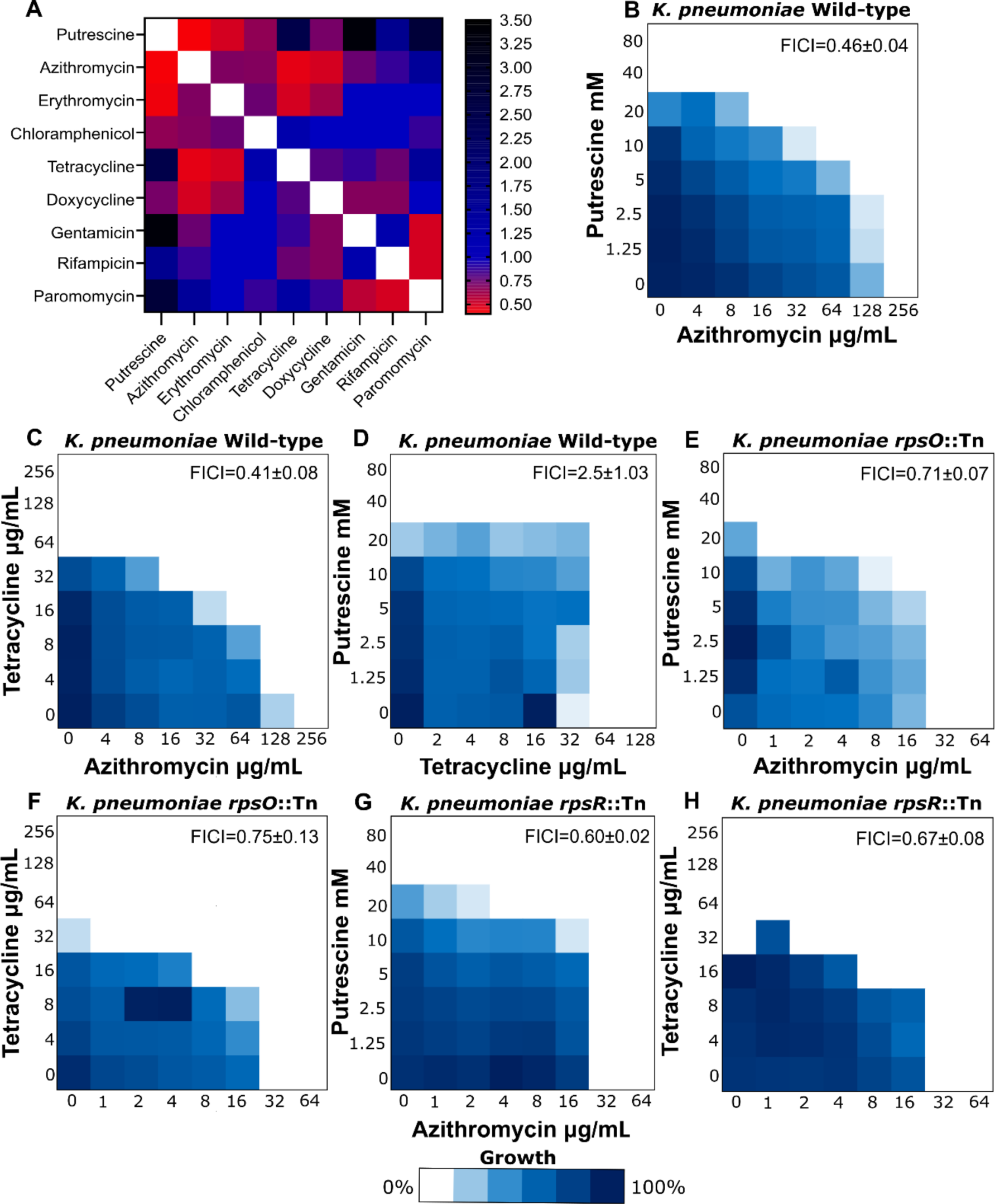
Putrescine may inhibit the ribosomal 30S subunit in a similar manner as tetracyclines. A) Heat map showing the average FICI value of duplicate checkerboard assays against *K. pneumoniae* MKP103 with various drug combinations. B-D) Checkerboard assays against *K. pneumoniae* (MKP103) treated with B) putrescine-azithromycin, C) tetracycline-azithromycin, and D) putrescine-tetracycline combinations. E-F) Checkerboard assays against *K. pneumoniae rpsO*::Tn treated with E) putrescine-azithromycin and F) tetracycline-azithromycin. G-H) Checkerboard assays against *K. pneumoniae rpsR*::Tn treated with G) putrescine-azithromycin and H) tetracycline-azithromycin. Dark blue regions represent high cell density determined by OD_600_. FICI mean and SEM, derived from 3 independent experiments, are displayed on each representative checkerboard.

Chloramphenicol binds to the aminoacyl site of the 50S ribosomal subunit and prevents peptidyl transferase. Azithromycin and erythromycin both block the 50S ribosomal subunit nascent peptide tunnel inhibiting transpeptidation. Tetracycline and doxycycline target the 30S ribosomal acceptor site preventing the association of aminoacyl-tRNA with the ribosome complex and prevent mRNA binding ^71^. We used rifampicin, an RNA polymerase inhibitor, as a control. Out of the seven antibiotics tested, only tetracyclines (tetracycline and doxycycline) synergized with azithromycin as putrescine did (Fig 5A). Synergy between the tetracycline doxycycline and azithromycin has been observed before in *P. aeruginosa*, albeit the underlying mechanism has not been resolved ^72^. Checkerboard assays of putrescine-azithromycin combinations were phenocopied by tetracycline-azithromycin combinations in the *K. pneumoniae* wild type strain (Fig 5B-C) and tetracycline also showed additive effects with putrescine (Fig 5D). Similarly, the additive effects of the putrescine-azithromycin combination against the ribosomal mutants *rpsO*::Tn and *rpsR*::Tn was also observed in the tetracycline-azithromycin combination (Fig 5E-H). Additionally, the combination of tetracycline and azithromycin, in checkerboards of PMBN treated *K. pneumoniae* (FICI 0.50 ± 0.00) and the *waaD*::Tn mutant still exhibited significant synergy (FICI 0.41 ± 0.07) (Fig S8A-C); under these conditions, while the outer membrane is perturbed chemically or genetically, synergy may still occur at the ribosomal level. Notably, with the exception of macrolides, the ribosome-targeting antibiotics we tested in Fig 5A are typically active against Gram-negative bacteria, in the absence of a specific resistance determinant; we further confirmed that the outer membrane does not serve as a barrier to their entry into *K. pneumoniae* MKP103 by showing a lack of MIC shift of these antibiotics between the wild-type strain and membrane permeable *waaD*::Tn mutant (Table S2). These results, along with the observations in the membrane permeable and ribosomal subunit mutants (Fig 5B-H and SA-C), suggest that the observed interactions between the tested ribosome-targeting antibiotics and putrescine or azithromycin – whether synergy or additive effects – are likely not driven by an outer membrane barrier effect but rather on-target effects on the ribosome.

Previously, *E. coli* serially passaged in sub-lethal putrescine concentrations evolved putrescine-resistant mutants that had truncations of the S7 30S ribosomal protein, a protein that organizes the folding of the 3’ major domain of 16S rRNA and crosslinks to the anticodon loop of tRNAs ^73, 74^. These mutants exhibited increased growth rates relative to the wild type when grown in the presence of putrescine ^74^. Indeed, adaptive resistance is likely to occur at the target for a compound, supporting our findings that putrescine may act on the 30S ribosomal subunit. Polyamine cross-linking in the 30S subunit leads to 16S rRNA structural changes ^68^; similarly, tetracycline has been shown to alter the structure of 16S rRNA ^75^. Moreover, 30S subunits lacking RpsO exhibit subtle changes in 16S rRNA nucleotide reactivity ^52, 76^. Thus, it is probable that the mutants of RpsO and RpsR lead to an altered 16S rRNA structure similar to that occurring as a result of polyamine and tetracycline binding to the 30S subunit, which may explain the lack of synergy of these drugs with azithromycin in *rpsO*::Tn and *rpsR*::Tn mutant backgrounds. Together, our results provide support that putrescine acts on the 30S ribosomal subunit similar to tetracycline and that it mediates the macrolide synergy via a dual mechanism (at the membrane and ribosomal levels).

### A screen of a polyamine analog library uncovers azithromycin synergists

We sought to exploit the polyamine-azithromycin synergy and the understanding of its mode of action to uncover an antibiotic adjuvant that may expand the spectrum of azithromycin to *K. pneumoniae*. We hypothesized that a synthetic polyamine analog may lead to synergy with azithromycin similar to that mediated by natural polyamines, serving as a potential macrolide adjuvant. To this end, we screened a library of 83 polyamine analogs against MKP103 alone and in combination with a sub-inhibitory concentration (1/8^th^ MIC) of azithromycin (Fig 6A). Six compounds displayed growth inhibition alone and in combination with azithromycin. Follow up checkerboard assays of the six primary hits revealed that OES2-017, OES2-046, OES2-045, and OES2-008 synergized with azithromycin at concentrations starting at the low micromolar range with FICI values of 0.54 ± 0.11, 0.32 ± 0.05, 0.37 ± 0.03, and 0.29 ± 0.05, respectively (Fig 6B-E). Similar to putrescine, the identified polyamine analogs showed growth inhibitory effects against *K. pneumoniae* MKP103; their MIC values (31.25 μM, 2.5, 2.5, and 5 mM for OES2-017, −045, −046, and −008, respectively, Fig 6B-E) were substantially lower than that of putrescine (40 mM).

**Figure 6.**
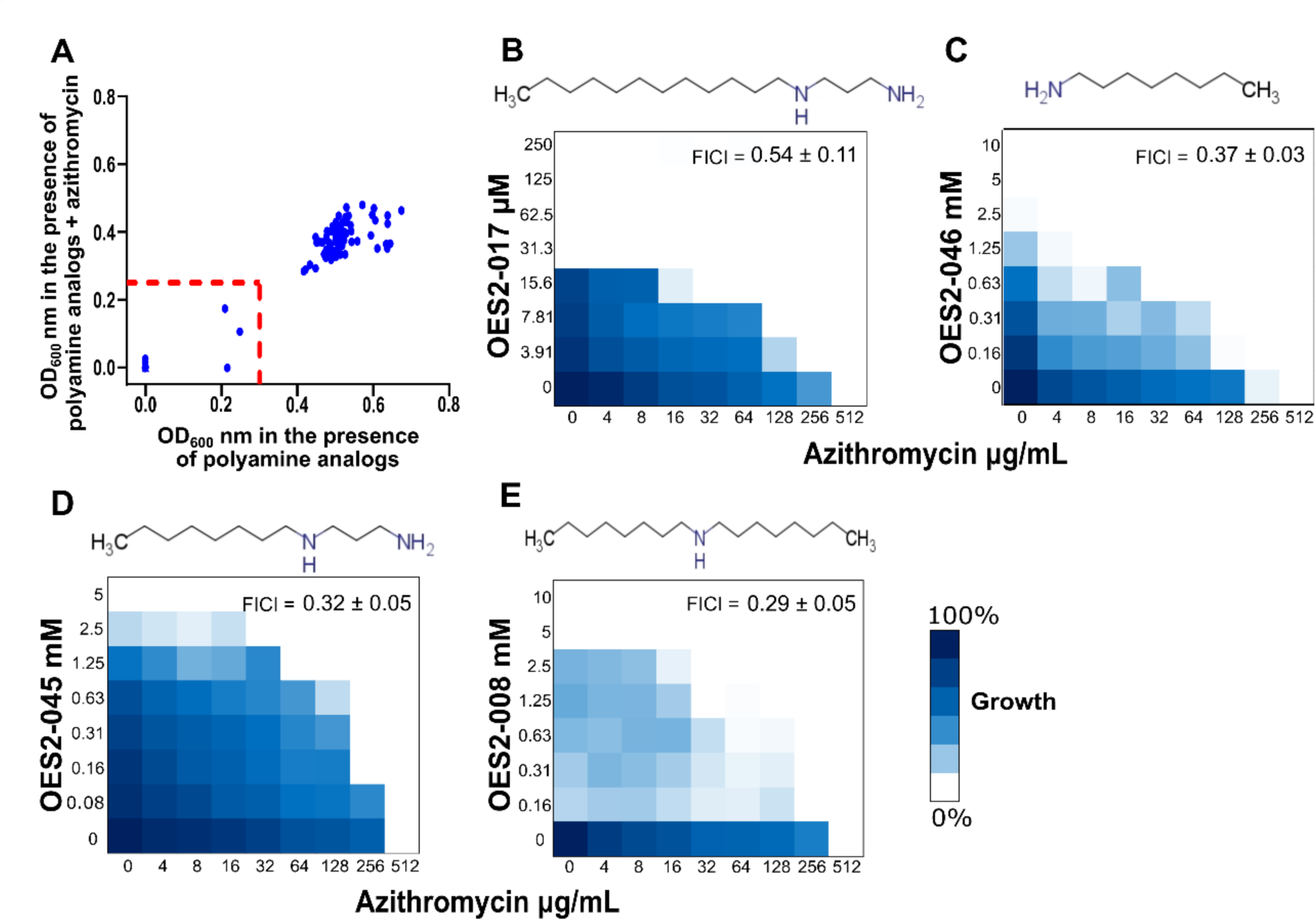
Screening for azithromycin adjuvants identifies polyamine analogs with more potent azithromycin synergy and growth inhibitory activities. A) A replica plot displaying the OD readings of *K. pneumoniae* MKP103 in the presence of the polyamine analogs at 2.5 mM alone (x-axis) and in combination with 64 µg/mL azithromycin (y-axis). B-E) Representative checkerboard assays of *K. pneumoniae* MKP103 treated with the prioritized polyamine analog hits in combination with azithromycin. All FICI values are shown as the average of at least 3 independent repeats and the SEM. Dark blue regions represent high cell density determined by OD_600_.

Next, we sought to identify the mechanism of azithromycin synergy by these four compounds, first by checking if they acted through a similar mechanism as that of putrescine. Therefore, we conducted checkerboard assays of each compound with azithromycin against the *rpsO*::Tn, *rpsR*::Tn, and *waaD*::Tn *K. pneumoniae* mutants, against which putrescine lost its synergy with the macrolide antibiotic, and tested their outer and inner membrane permeabilization effects. Both OES2-017 and OES2-008 showed reduced or complete loss of azithromycin synergy against all three mutants (Fig S9A-F). Conversely, OES2-045 and OES2-046 exhibited no loss of synergy against the mutants, except that the extent of the synergy against *waaD*::Tn (mean FICI= 0.419 and 0.489, respectively) was reduced relative to that against the wild type (mean FICI= 0.325 and 0.375, respectively) (Fig S9G-L). We showed in NPN fluorometric kinetic assays that OES2-017, OES2-008, OES2-045, and OES2-046, in descending order of magnitude, disrupted the outer membrane of *K. pneumoniae* in a dose dependant manner across the synergistic range of each compound (Fig S10A-D). Further, all four analogs exhibited inner membrane activity, in colorimetric ONPG hydrolysis assays against *E. coli* ML-35, in a dose dependant manner across the synergistic range of each compound (Fig S11A-D). Taken together, these results suggest that OES2-017 and OES2-008 synergize with azithromycin via mechanisms similar to those of putrescine at the ribosome and outer and inner membranes levels, whereas OES2-045 and OES2-046 may exert their azithromycin synergy through membrane activity only. Importantly, these analogs synergize with azithromycin at substantially lower concentrations than putrescine, serving as potential macrolide adjuvants.

## Conclusions

In this study, we uncovered a sensitization of *K. pneumoniae* to the macrolide azithromycin in the presence of potentially physiological levels of polyamines. We showed that the putrescine-azithromycin synergy occurs in pan-drug-resistant and hypervirulent *K. pneumoniae* lineages, bringing azithromycin susceptibility in 78 of 100 clinical isolates below the CLSI breakpoint. Further, we have provided evidence towards a dual mechanism of action underlying the putrescine-azithromycin synergy, whereby the polyamine: 1. permeabilized the outer and inner membranes to azithromycin and 2. compounded the protein synthesis inhibiting effects of azithromycin by acting at the ribosome, likely at the 30S subunit (Fig 7).

**Figure 7.**
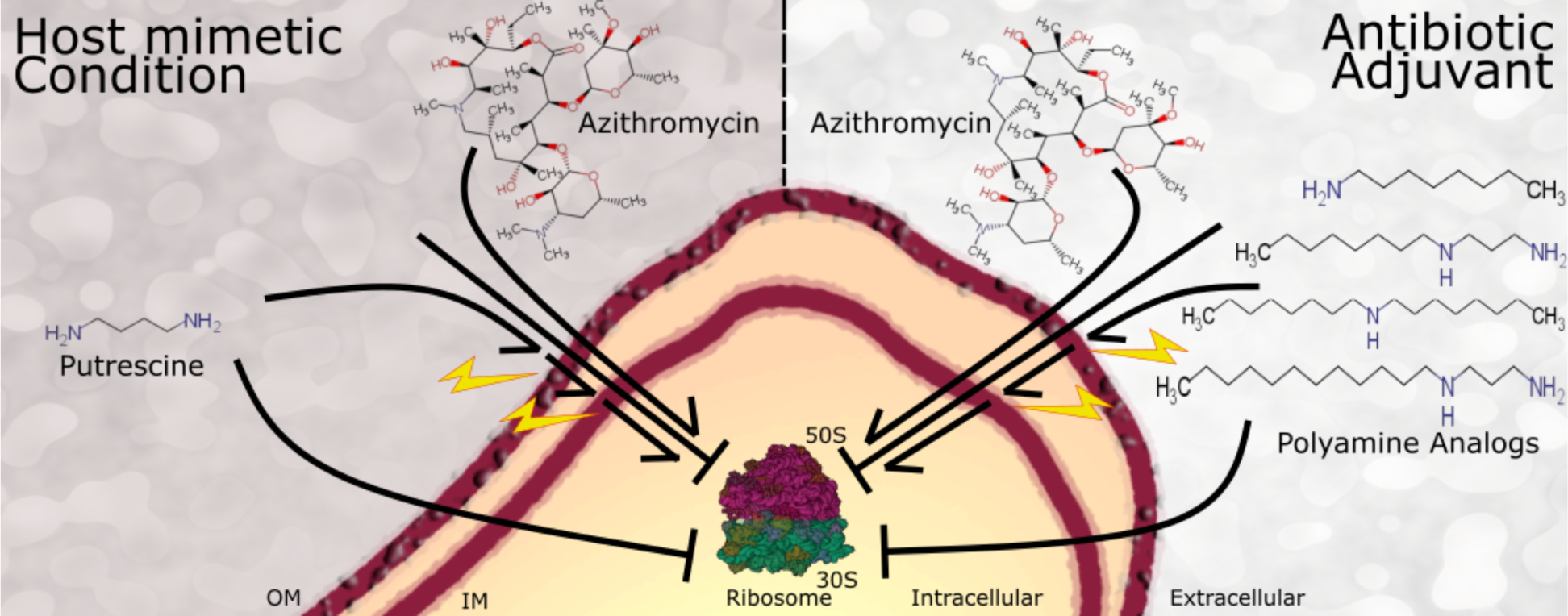
A model of the proposed dual mode of action of polyamine-azithromycin synergy. The natural polyamine, putrescine, and the synthetic polyamine analogs identified herein synergize with azithromycin via: 1. Perturbation of both the outer and inner membranes of Gram-negative bacteria and 2. Compounding azithromycin protein synthesis inhibition whereby azithromycin inhibits the 50S ribosomal subunit and putrescine likely acts on the 30S subunit of the ribosome.

The exact physiological levels of polyamines that bacteria may encounter during infection is not known. Polyamine levels vary across different body sites as shown in a mouse model experiment revealing organ tissue polyamine levels different across the skin, heart, liver, and kidney^38^. The same appears to be true in humans. For example, putrescine concentration was determined to be 3 mM in urine ^77^. Other studies showed that putrescine reached up to 0.2 mM in sputum samples from CF patients in other studies ^35, 78^; however, infection alters the rheology of the mucus and the lung environment precluding accurate prediction of the local concentration of putrescine and other polyamines in the lung of CF patients^79^. Moreover, putrescine levels increase dramatically (by ≥8-fold) during exacerbations of bacterial infections in CF patients ^35,78^. Further, the majority of intracellular polyamines in eukaryotic cells (≥85%) are typically found bound to macromolecules, including DNA, RNA, and phospholipids ^80, 81^; thus, the above estimates of free polyamines might represent an underestimation of the total intracellular polyamine pools. Hence, the concentrations used in this study could potentially resemble the physiological cumulative concentrations of all polyamines bacteria may encounter during infection in certain body compartments.

We aimed to uncover new antibiotic adjuvants that mimic the mechanism natural polyamines synergize with azithromycin to circumvent the uncertainty of the level of host polyamines by acting in a controlled manner and ideally at significantly lower concentrations than the natural polyamines. To that end, we screened a library of polyamine analogs and discovered four synthetic polyamine analogs that displayed growth inhibitory effects and synergy with azithromycin against *K. pneumoniae*, with greater potency compared to natural polyamines (Fig 7). Synergy with currently available Gram-positive specific antibiotics, such as macrolides, including by perturbing the Gram-negative outer membrane, represents a potential strategy to combat drug-resistant Gram-negative bacteria ^59^. As such, the polyamine analogs discovered herein may broaden the spectrum of azithromycin, serving as a potential strategy to combat antimicrobial resistant *K. pneumoniae*.

## Methods

### Strains, reagents, and culture conditions

Table 1 lists bacteria and plasmids used in this work. Bacteria were grown in cation-adjusted Mueller Hinton medium (CAMHB, herein referred to as MHB) at 37 °C buffered with 100 mM Tris-HCl pH7.4 buffer (to offset the alkalinity of higher polyamine concentrations), unless otherwise indicated. Antibiotics and reagents were obtained from Sigma unless otherwise indicated.

**Table 1.**
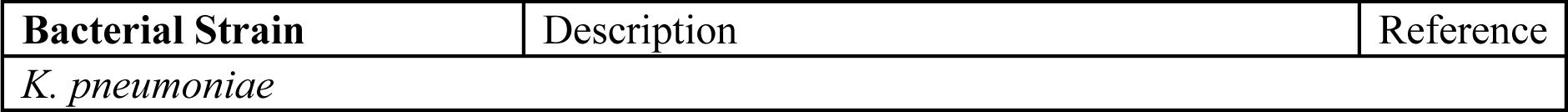
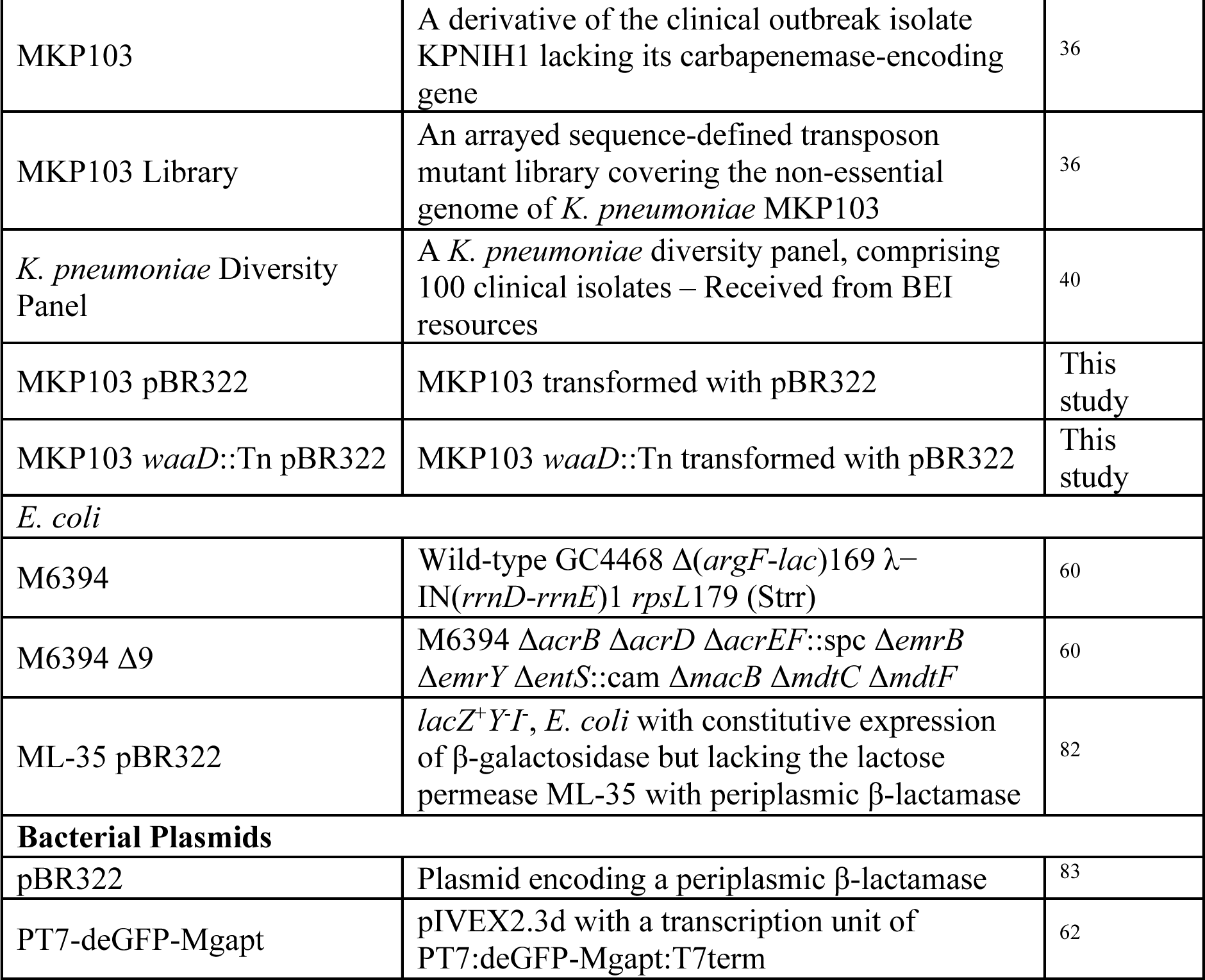
Bacterial strains and plasmids used in this study.

| Bacterial Strain | Description | Reference |
| --- | --- | --- |
| *K. pneumoniae* |  |  |
| MKP103 | A derivative of the clinical outbreak isolate KPNIH1 lacking its carbapenemase-encoding gene | 36 |
| MKP103 Library | An arrayed sequence-defined transposon mutant library covering the non-essential genome of *K. pneumoniae* MKP103 | 36 |
| *K. pneumoniae* Diversity Panel | A *K. pneumoniae* diversity panel, comprising 100 clinical isolates – Received from BEI resources | 40 |
| MKP103 pBR322 | MKP103 transformed with pBR322 | This study |
| MKP103 *waaD*::Tn pBR322 | MKP103 *waaD*::Tn transformed with pBR322 | This study |
| *E. coli* |  |  |
| M6394 | Wild-type GC4468 $\Delta(\arg F-lac)169 \lambda^{-}$ IN(*rrnD-rrnE*)1 *rpsL*179 (Strr) | 60 |
| M6394 $\Delta 9$ | M6394 $\Delta acrB \Delta acrD \Delta acrEF::spc \Delta emrB \Delta emrY \Delta entS::cam \Delta macB \Delta mdtC \Delta mdtF$ | 60 |
| ML-35 pBR322 | *lacZ*<sup>+</sup>*Y-I*, *E. coli* with constitutive expression of β-galactosidase but lacking the lactose permease ML-35 with periplasmic β-lactamase | 82 |
| **Bacterial Plasmids** |  |  |
| pBR322 | Plasmid encoding a periplasmic β-lactamase | 83 |
| PT7-deGFP-Mgapt | pIVEX2.3d with a transcription unit of PT7:deGFP-Mgapt:T7term | 62 |

### Antimicrobial susceptibility testing

All MIC and checkerboard assays were conducted following the CLSI method for MIC testing by broth microdilution ^84^. The assays were conducted in polystyrene 96-well plates with final volume of 100 μL/well and inoculum size adjusted as optical density at 600 nm (OD_600_) of 0.001. Plates were incubated while shaking at 600 RPM and 37°C; growth of the bacterial culture was determined by OD_600_ measurement at 24 hours. To assess the combined antimicrobial activity, fractional inhibitory concentration indices (FICI) were calculated as FICI = A/MIC_A_ + B/MIC_B_. A and B represent the MIC of the drugs while in combination. MIC_A_ and MIC_B_ represent the individual MIC of each drug alone. FICI values less than or equal to 0.5 were considered synergism, values greater than 4 were considered antagonism, and values between 0.5 and 4 were indicated indifference, additive, or no interaction.

### Arrayed transposon *Klebsiella pneumoniae* MKP103 library screen

Overnight cultures of the *K. pneumoniae* MKP103 library^36^ were performed using the BioMatrix benchtop robot (S&P robotics) into 384 well-density plates containing MHB with 150 μg/mL of chloramphenicol and incubated at 37°C for 24 hours. Subsequently, the entire library was inoculated into MHB with azithromycin at 32 μg/mL and 64 μg/mL and without azithromycin. The plates were incubated at 37°C, and OD_600_ was read after 24 hours.

### NPN assays

The outer membrane permeability of *K. pneumoniae* MKP103 and *E. coli* M6394 wild type and mutants was measured using NPN uptake assays as previously described ^85^.

Briefly, bacteria were subcultured in MHB until mid to late log phase, washed, and resuspended in either 100 mM Tris-HCl (pH 7.4) with 20 mM glucose or 5 mM HEPES (pH7.4) with 20 mM glucose. The latter buffer was used only when low putrescine concentrations, causing minimal pH changes, were tested. NPN was added to the bacterial suspension at a final concentration of 20 µM and dispensed into black 96 well plates pre-seeded with a concentration gradient of putrescine to a total final volume of 200 µL. We tested untreated bacteria suspended in the same buffer adjusted to the final pH reached upon addition of each tested putrescine concentration as a negative control (untreated pH control) used to correct for any pH related signal. Fluorescence was read kinetically (excitation λ =355 nm, emission λ =420 nm) every 5 minutes for 215 minutes. The %NPN uptake was calculated using the following equation: %NPN uptake = (F_obs_-F_0_)/(F_100_-F_0_) x 100, where F_obs_ is the observed fluorescence at a given concentration of putrescine, F_0_ is the fluorescence in non-treated cells (untreated pH control), and F_100_ is the fluorescence of NPN in 8 µg/mL colistin-treated cells (used as a positive control).

### β-lactamase assay

The outer membrane integrity of *K. pneumoniae* wild type and *waaD*::Tn mutant were tested by measuring β-lactamase activity with nitrocefin hydrolysis as previously described ^86^. First, we extracted pBR322 from an *E. coli* strain carrying the plasmid (Monarch Mini-Prep kit, NEB) and transformed it into electrocompetent *K. pneumoniae* strains of interest by electroporation^36^ Overnight cultures of *K. pneumoniae* pBR322 and *K. pneumoniae waaD*::Tn pBR322 were diluted 100-fold into TSB and incubated at 37°C until late log phase (OD_600_ of ∼1.8). Aliquots of the sub-cultures were centrifuged and washed twice with PBS then resuspended in 100 mM Tris-HCl buffer (pH 7.4) at a final OD_600_ of 0.01. The bacterial suspensions mixed with 30 μM nitrocefin were dispensed into 96 well plates pre-seeded with the tested drugs with a total final volume of 100 μL/well. The plates were incubated at 37°C while measuring the absorbance kinetically at 492 nm every 10 minutes.

### β-galactosidase assay

The inner membrane integrity of *E. coli* ML-35 pBR322 was tested by measuring β-galactosidase activity with ONPG hydrolysis as previously described ^86^. An overnight culture of *E. coli* ML-35 pBR322 was diluted 100-fold into TSB and incubated at 37°C until late log phase (OD_600_ of ∼1.8). Aliquots of the sub-culture were centrifuged and washed twice with PBS then resuspended in 100 mM Tris-HCl buffer (pH 7.4) at a final OD_600_ of 0.01. The bacterial suspensions mixed with 1.5 mM ONPG were dispensed into 96 well plates pre-seeded with the tested drugs with a total final volume of 100 μL/well. The plates were incubated at 37°C while measuring the absorbance kinetically at 405 nm every 2 minutes.

### Cell free protein expression

An NEB Express Cell-free *E. coli* protein synthesis system (New England Biolabs) was used for *in vitro* protein expression. Transcription and translation were measured fluorometrically (excitation/emission wavelengths at 610/650 nm MGapt, 485/525 nm deGFP) using an engineered plasmid that encodes a deGFP protein and a malachite green RNA aptamer (PT7-deGFP-MGapt was a gift from Richard Murray, via Addgene plasmid # 67741; http://n2t.net/addgene:67741) ^62^. The cell-free protein synthesis system reaction mixture was prepared following the manufacturer’s protocol and dispensed as 5 µL aliquots into black 384 well plates. Putrescine, dH_2_O, plasmid (124 ng/µL final concentration), and Malachite green dye (10 µM final concentration) top up the reaction mixture reaching a combined final volume of 11 µL.

### Polyamine analog library screen

*K. pneumoniae* MKP103 was inoculated in MHB with 100 mM Tris-HCl buffer (pH 7.4) at a 0.001 OD_600_ and dispensed into 96 well plates at a total final volume of 100 μL (including polyamine analogs, azithromycin, vehicle, or combinations thereof). Polyamine analogs were screened at 2.5 mM alone and in combination with 64 µg/mL azithromycin. Plates were incubated at 37°C and 600 RPM; growth of the bacterial culture was determined by OD_600_ readings at 24 hours.

### Statistical analysis

All statistical analyses were calculated using GraphPad Prism 9.

### Supporting Information Available free of charge

Fig S1 An index plot of the screen of *K. pneumoniae* MKP103 mutants grown in 64 μg/mL of azithromycin; Fig S2 Putrescine-azithromycin checkerboard assays of mutants of putative determinants from the chemogenomic screen; Fig S3 LPS mutants, their LPS structure, and their resulting putrescine-azithromycin checkerboards; Fig S4 Confirmation of deep rough LPS mutation in *waaD*::Tn mutant; Fig S5 Genetically perturbed outer membrane leads to a loss of putrescine-azithromycin synergy; Fig S6 *K. pneumoniae* NPN assay showing that low putrescine concentrations stabilize the outer membrane of *K. pneumoniae* MKP103; Fig S7 Cell-free fluorometric mRNA and protein synthesis assay showing that putrescine inhibits protein synthesis; Fig S8 Tetracycline-azithromycin checkerboard assays of membrane-perturbed *K. pneumoniae* MKP103; Fig S9 Checkerboard assays of *K. pneumoniae* MKP103 mutants treated with synthetic polyamine analogs and azithromycin; Fig S10 Polyamine analog outer membrane perturbation NPN assays in *K. pneumoniae* MKP103; Fig S11 Polyamine analog inner membrane perturbation ONPG assays in *E. coli* ML-35 pBR322; Table S1 Putrescine-azithromycin FICI values of *K. pneumoniae* diversity panel isolates; Table S2 MIC comparison of ribosomal targeting antibiotics in *K. pneumoniae* wild-type and *waaD*::Tn.

## Acknowledgments.

This work was funded by a Natural Sciences and Engineering Research Council of Canada discovery grant, a Saskatchewan Health Research Foundation establishment grant, and startup funds from the Faculty of Science at the University of Regina. O.M.E. holds a Canada Research Chair in Chemogenomics and Antimicrobial Research. P.B.M. was supported by a CIHR Canada Graduate Scholarship – Master’s (CGS-M) award.

