## Supplementary figures and tables for "Polyamine-mediated sensitization of *Klebsiella pneumoniae* to macrolides through a dual mode of action"

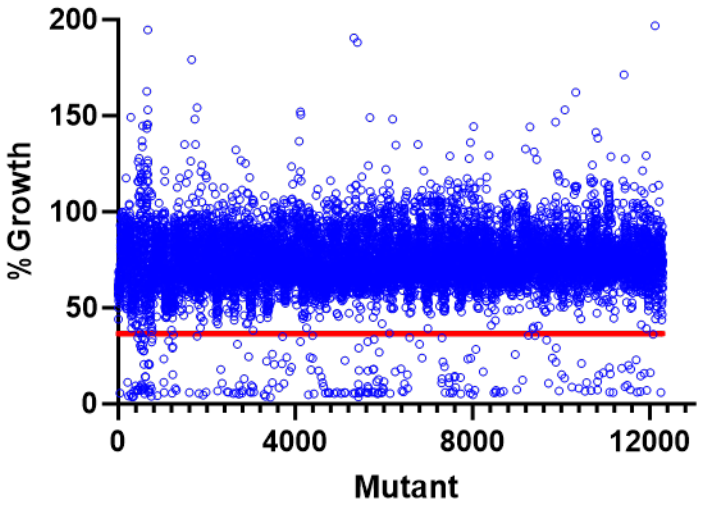


**Figure S1.** An index plot of the screen of *K. pneumoniae* MKP103 mutants grown in 64 µg/mL of azithromycin shown as % growth to that of the untreated mutants. The red line denotes the cutoff under which datapoints were considered putative hits.

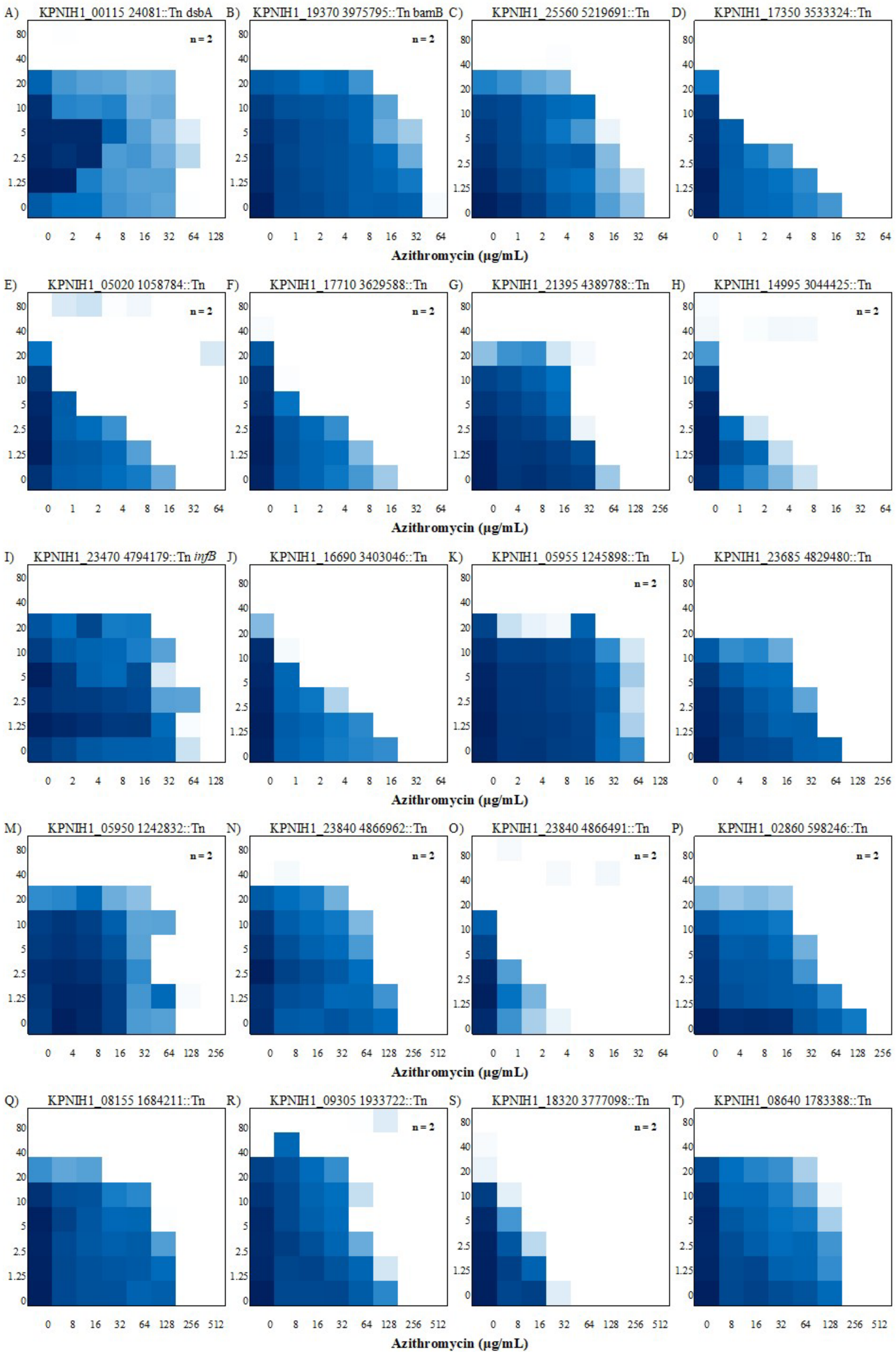
A) KPNIH1_00115 24081::Tn dsbA
B) KPNIH1_19370 3975795::Tn bamB
C) KPNIH1_25560 5219691::Tn
D) KPNIH1_17350 3533324::Tn
E) KPNIH1_05020 1058784::Tn
F) KPNIH1_17710 3629588::Tn
G) KPNIH1_21395 4389788::Tn
H) KPNIH1_14995 3044425::Tn
I) KPNIH1_23470 4794179::Tn infB
J) KPNIH1_16690 3403046::Tn
K) KPNIH1_05955 1245898::Tn
L) KPNIH1_23685 4829480::Tn
M) KPNIH1_05950 1242832::Tn
N) KPNIH1_23840 4866962::Tn
O) KPNIH1_23840 4866491::Tn
P) KPNIH1_02860 598246::Tn
Q) KPNIH1_08155 1684211::Tn
R) KPNIH1_09305 1933722::Tn
S) KPNIH1_18320 3777098::Tn
T) KPNIH1_08640 1783388::Tn
n = 2
Azithromycin (µg/mL)

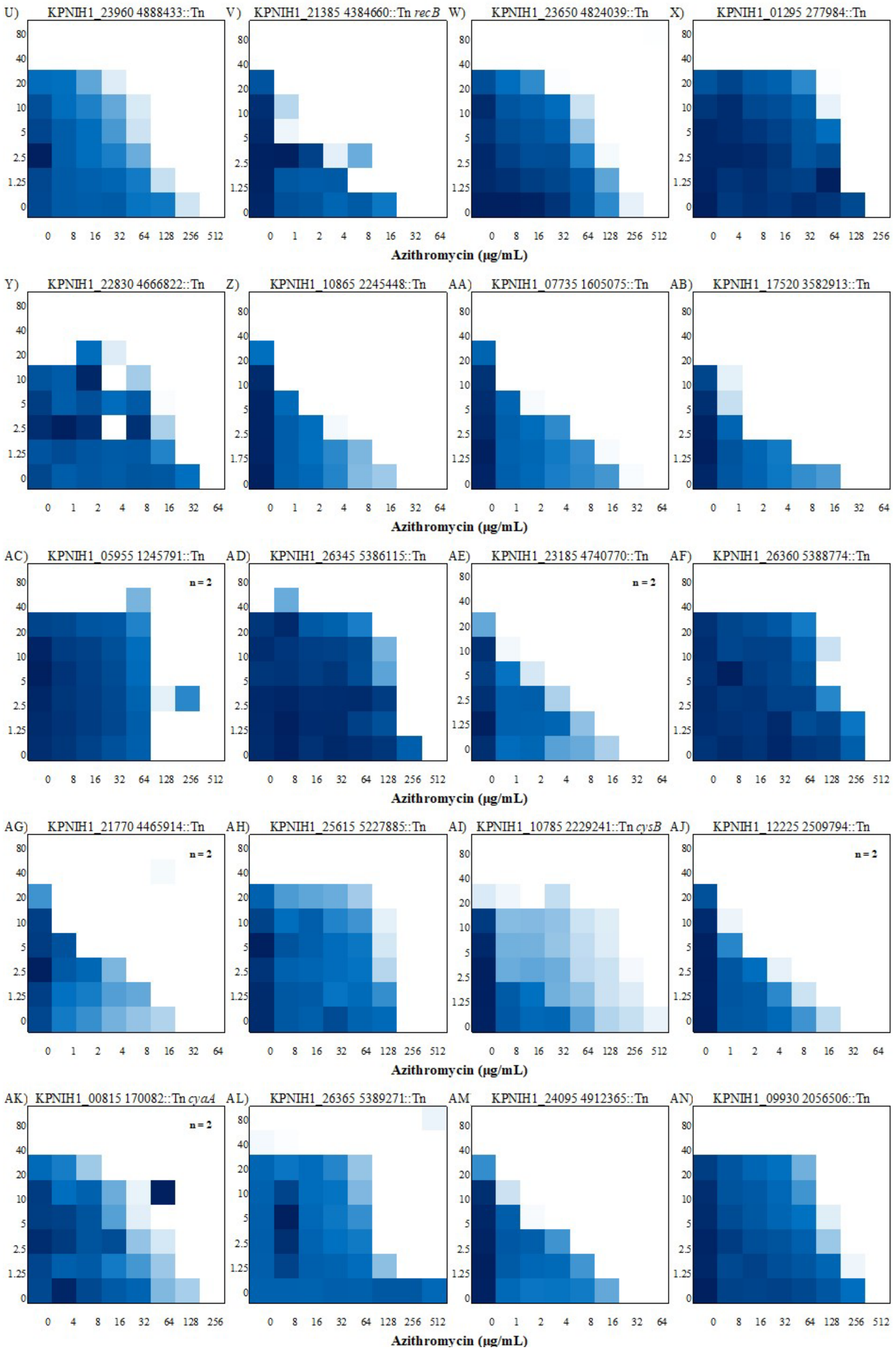
U) KPNIH1_23960 4888433::Tn
V) KPNIH1_21385 4384660::Tn *recB*
W) KPNIH1_23650 4824039::Tn
X) KPNIH1_01295 277984::Tn
Y) KPNIH1_22830 4666822::Tn
Z) KPNIH1_10865 2245448::Tn
AA) KPNIH1_07735 1605075::Tn
AB) KPNIH1_17520 3582913::Tn
AC) KPNIH1_05955 1245791::Tn
n = 2
AD) KPNIH1_26345 5386115::Tn
AE) KPNIH1_23185 4740770::Tn
n = 2
AF) KPNIH1_26360 5388774::Tn
AG) KPNIH1_21770 4465914::Tn
n = 2
AH) KPNIH1_25615 5227885::Tn
AI) KPNIH1_10785 2229241::Tn *cysB*
AJ) KPNIH1_12225 2509794::Tn
n = 2
AK) KPNIH1_00815 170082::Tn *cyaA*
n = 2
AL) KPNIH1_26365 5389271::Tn
AM KPNIH1_24095 4912365::Tn
AN) KPNIH1_09930 2056506::Tn
Azithromycin (μg/mL)

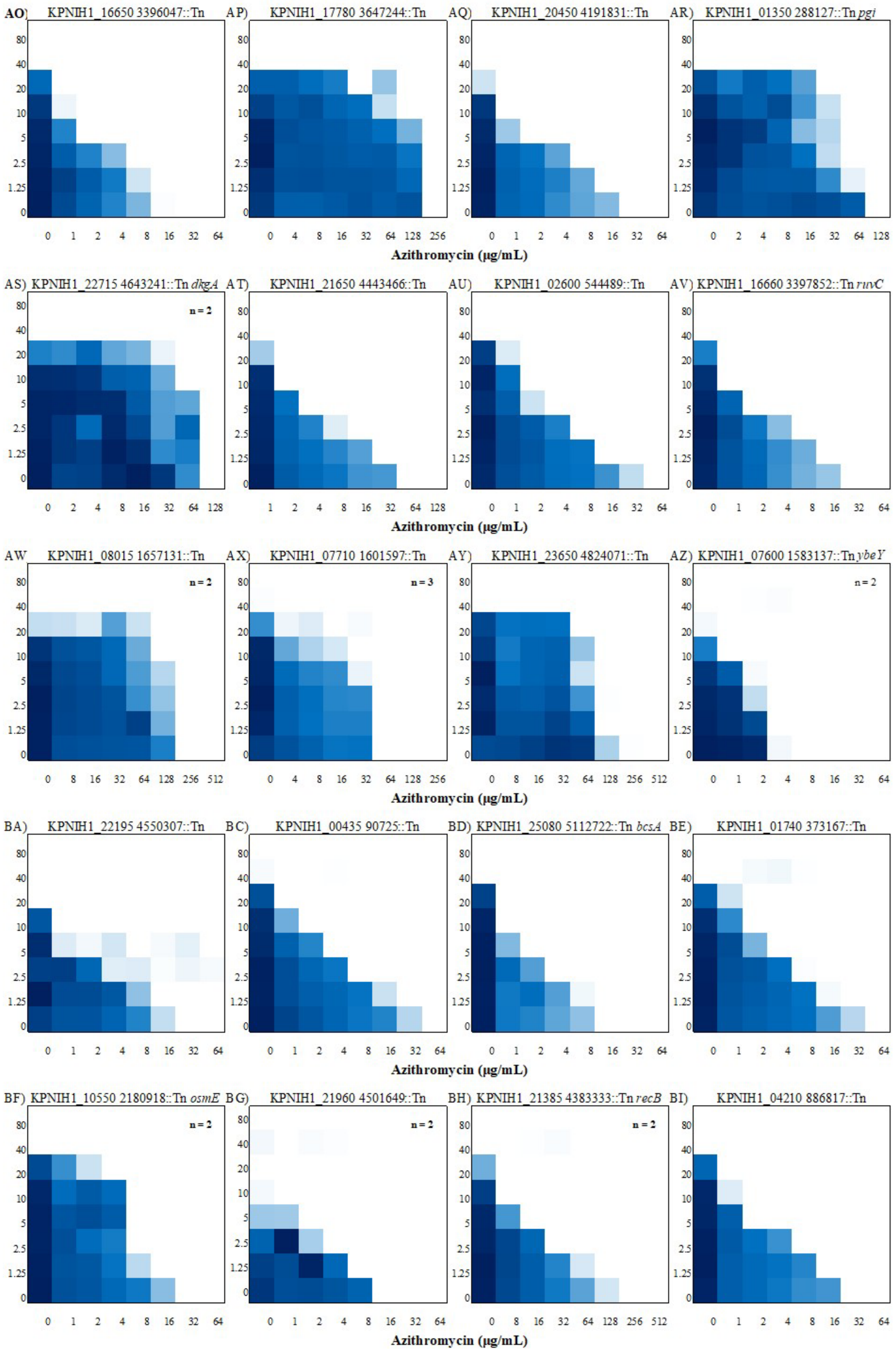
AO) KPNIH1_16650 3396047::Tn
AP) KPNIH1_17780 3647244::Tn
AQ) KPNIH1_20450 4191831::Tn
AR) KPNIH1_01350 288127::Tn pgi
AS) KPNIH1_22715 4643241::Tn dkgA
AT) KPNIH1_21650 4443466::Tn
AU) KPNIH1_02600 544489::Tn
AV) KPNIH1_16660 3397852::Tn ruvC
AW KPNIH1_08015 1657131::Tn
AX) KPNIH1_07710 1601597::Tn
AY) KPNIH1_23650 4824071::Tn
AZ) KPNIH1_07600 1583137::Tn ybeY
BA) KPNIH1_22195 4550307::Tn
BC) KPNIH1_00435 90725::Tn
BD) KPNIH1_25080 5112722::Tn bcsA
BE) KPNIH1_01740 373167::Tn
BF) KPNIH1_10550 2180918::Tn osmE
BG) KPNIH1_21960 4501649::Tn
BH) KPNIH1_21385 4383333::Tn recB
BI) KPNIH1_04210 886817::Tn
n = 2
n = 3
Azithromycin (μg/mL)

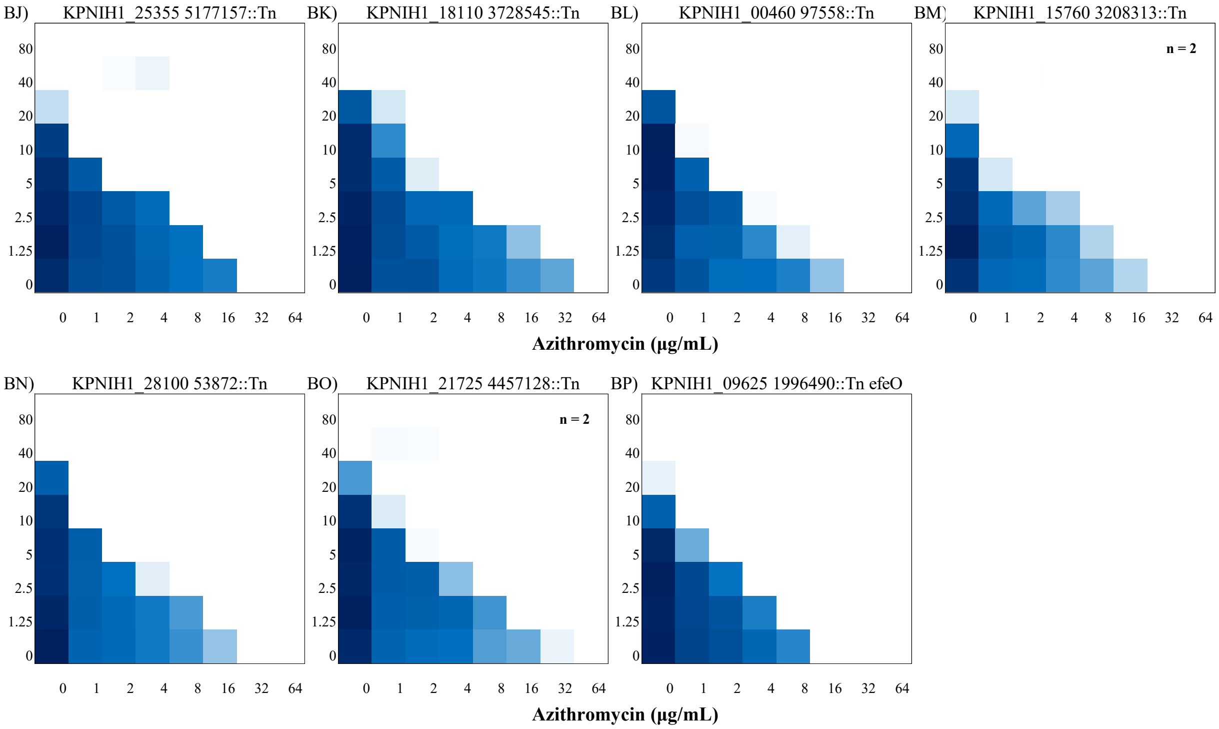


**Figure S2. Putrescine-azithromycin checkerboard assays of mutants of putative determinants from the chemogenomic screen.** Heat maps of representative putrescine (y-axis) – azithromycin (x-axis) checkerboards of the *K. pneumoniae* MKP103 mutants identified from the chemogenomic screen. (n) denotes number of replicates completed for mutants with more than one replicate. Dark blue represents maximal growth determined by $OD_{600}$.

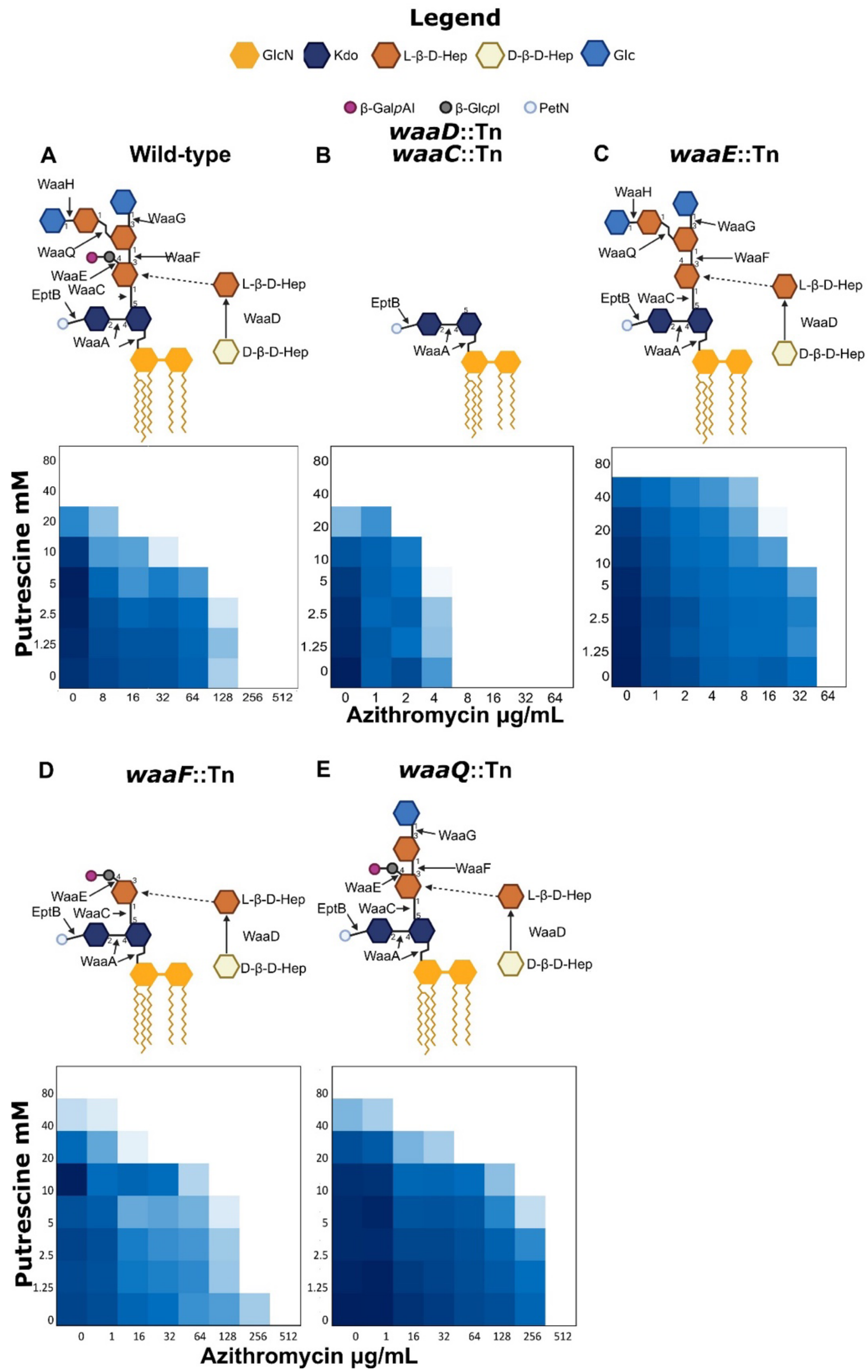


**Figure S3. LPS mutants, their LPS structure, and their resulting putrescine-azithromycin checkerboards.** Heat maps of representative putrescine-azithromycin checkerboards of the *K. pneumoniae* MKP103 LPS mutants. All checkerboards are representative of triplicate assays. Panel B shows a representative *waaD*::Tn checkerboard, Fig 2 shows representative *waaD*::Tn and *waaC*:Tn checkerboards. LPS structures shown are as predicted based on KEGG gene annotation of KPNIH10 strain[1, 2]. Dark blue represents maximal growth determined by $OD_{600}$.

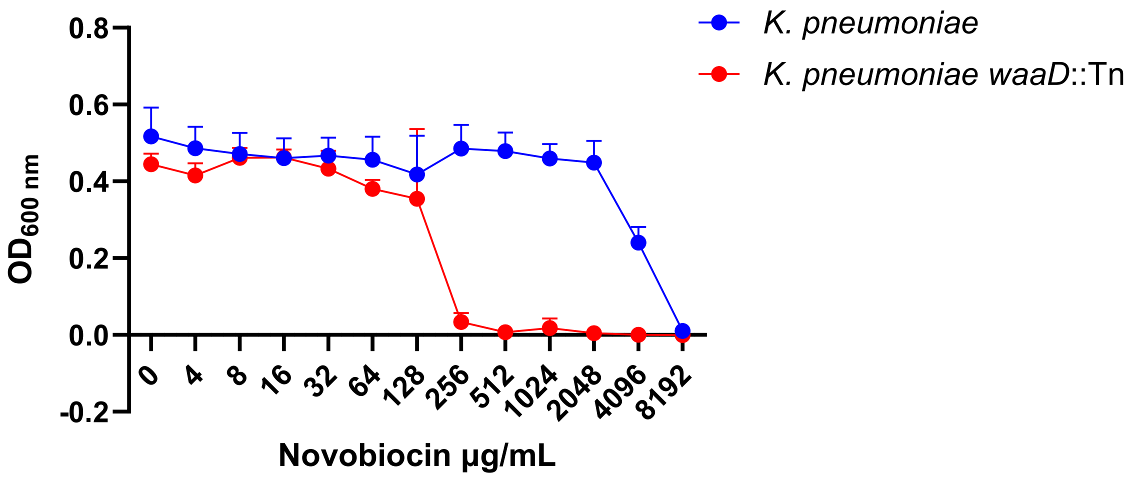


**Figure S4. Confirmation of deep rough LPS mutation in *waaD*::Tn mutant.** An MIC of *K. pneumoniae* MKP103 wild type (blue) and *waaD*::Tn mutant (red). The MIC graph represents the combined data of two independent MIC experiments each run in duplicate. Error shown is SEM.

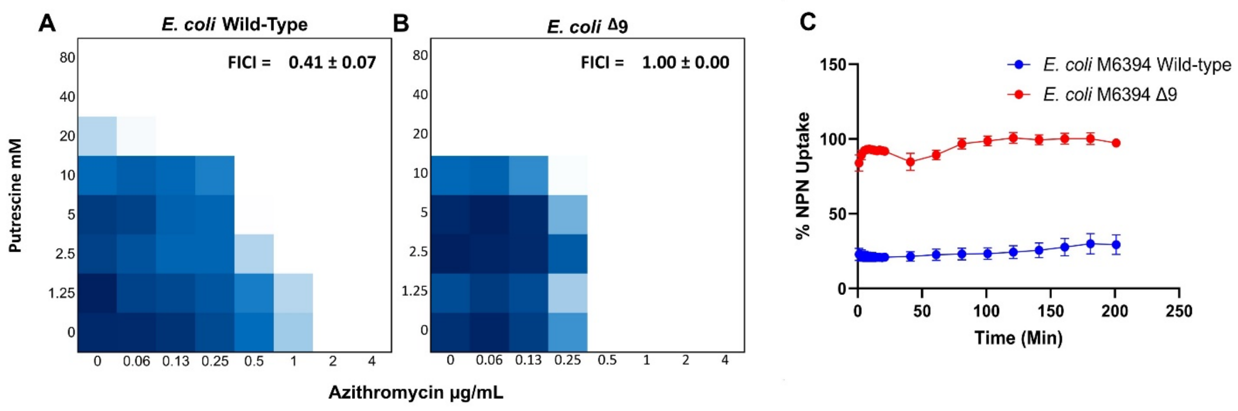


**Figure S5. Genetically perturbed outer membrane leads to a loss of putrescine-azithromycin synergy.** Representative of three independent putrescine-azithromycin checkerboard assays of A) *E. coli M6394* wild type and B) *E. coli* M6394 Δ9 (*ΔacrB ΔacrD ΔacrEF::spc ΔemrB ΔemrY ΔentS::cam ΔmacB ΔmdtC ΔmdtF*). Checkerboard assays were conducted in unbuffered cation-adjusted MHB. Dark blue represents maximal growth determined by $OD_{600}$. C) A fluorometric kinetic assay displaying grouped data from two independent experiments (n=6) of % NPN uptake in *E. coli* M6394 wild type and *E. coli* M6394 Δ9. Error shown as SEM.

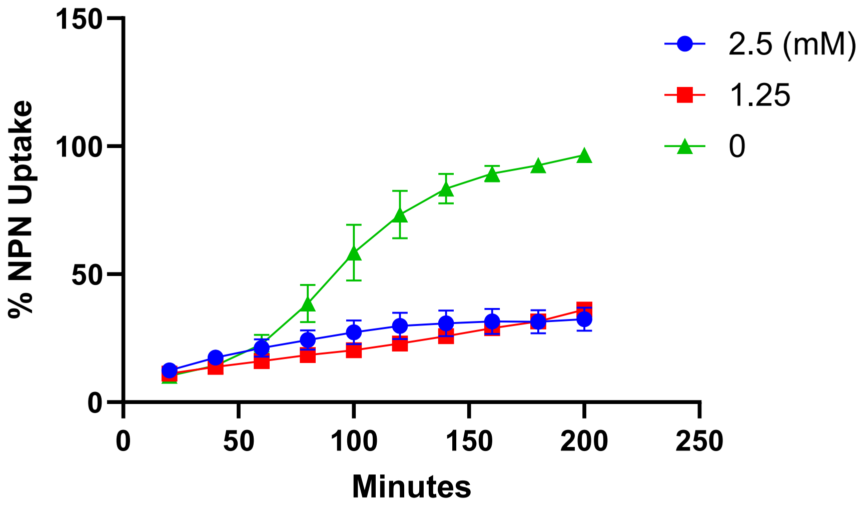


**Figure S6. Low putrescine concentrations stabilize the outer membrane of *K. pneumoniae* MKP103.** A fluorometric kinetic assay displaying grouped data from two independent experiments (n=6) of % NPN uptake in *K. pneumoniae* MKP103 suspended in 5 mM HEPES buffer (pH 7.4) and 20 mM Glucose. Error shown is SEM.

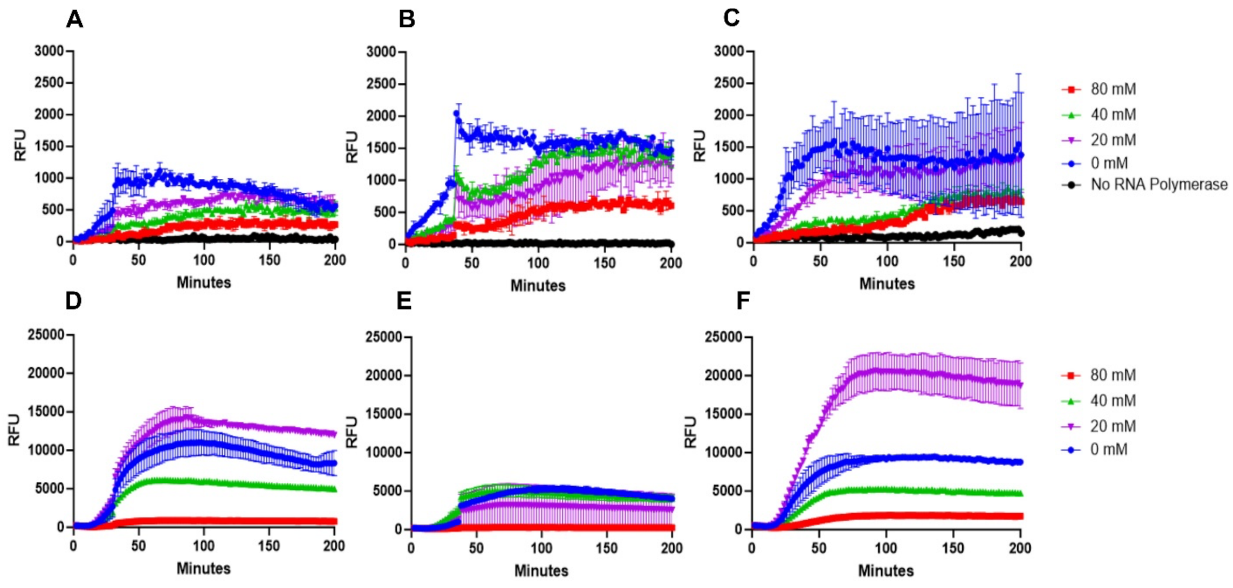


**Figure S7. Putrescine inhibits protein synthesis**. Kinetic reads of fluorometric experiments simultaneously tracking A-C) mRNA levels via RFU of mRNA-MG aptamer (610/650 nm) and D-F) Protein synthesis via RFU of deGFP expression (433/475 nm) in black 384 well plates using the NEBExpress cell-free protein expression system in 200 mM HEPES buffer with an engineered plasmid encoding deGFP with a malachite green mRNA aptamer. Each of the following graph pairs (A, D), (B, E), and (C, F) represent the corresponding mRNA and deGFP signals, respectively, from the same experiment. Error shown is SEM.

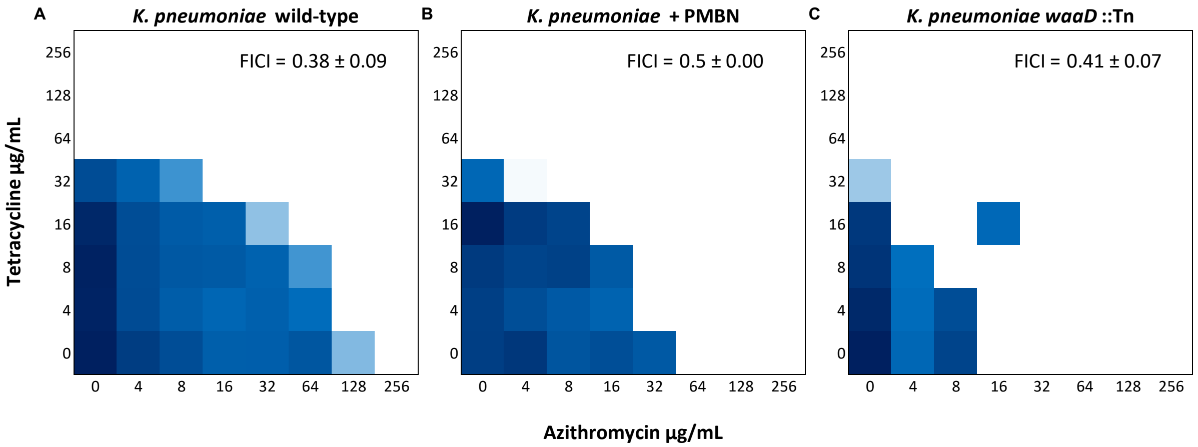


**Figure S8.** Heat maps of representative tetracycline-azithromycin checkerboards of *K. pneumoniae* MKP103 (n=2) in the A) absence and B) presence of 1 µg/mL PMBN. C) A checkerboard of *K. pneumoniae waaD*::Tn treated with tetracycline-azithromycin. Dark blue represents maximal growth determined by $OD_{600}$; error shown is SEM.

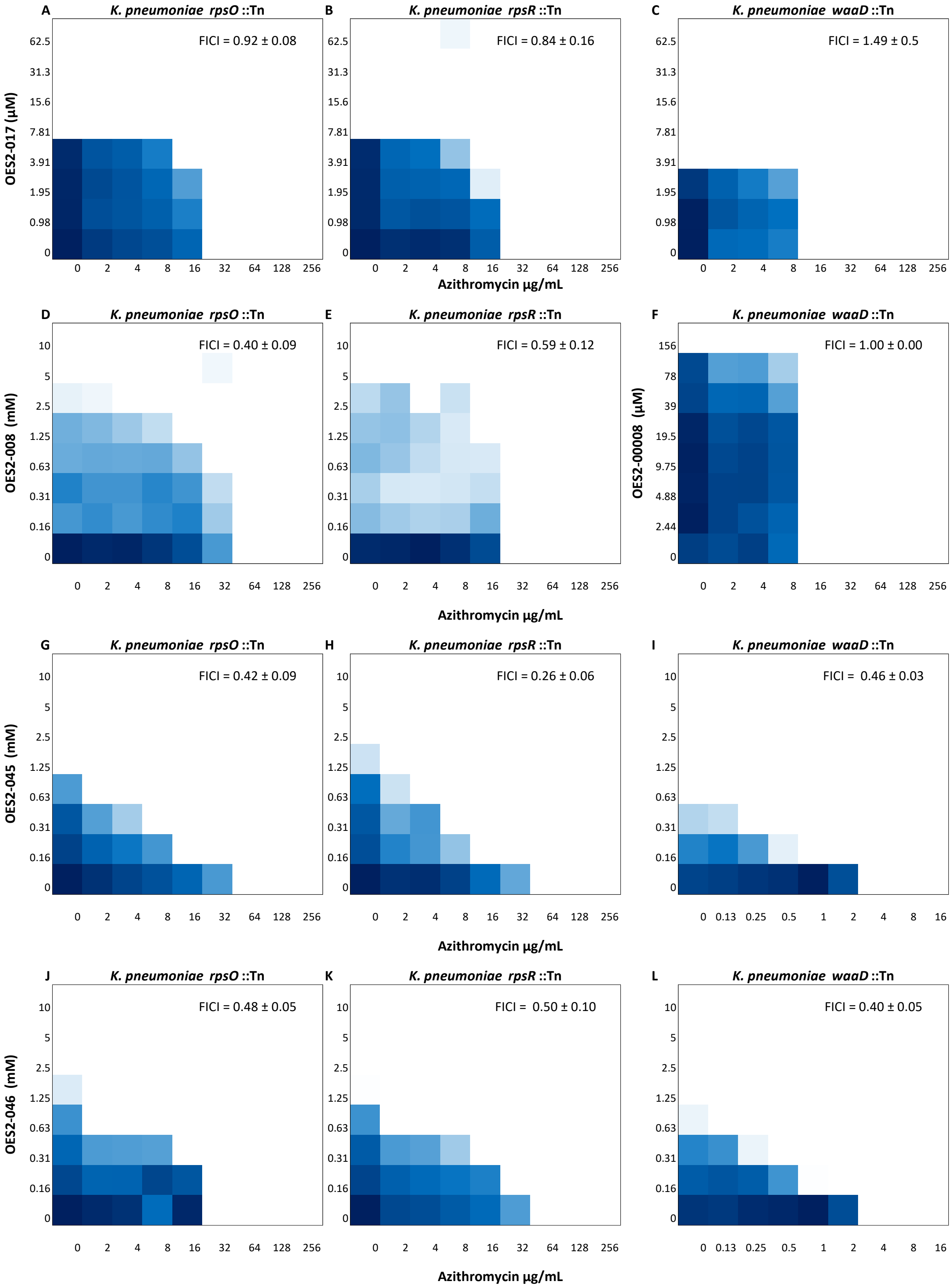


**Figure S9.** Heat maps of representative checkerboards of *K. pneumoniae* MKP103 mutants *rpsO*::Tn, *rpsR*::Tn, and *waaD*::Tn conducted in triplicate, treated with the polyamine analogs (OES2-017, OES2-008, OES2-045, OES2-046) on the y-axis and azithromycin (x-axis). Dark blue represents maximal growth determined by $OD_{600}$ (error shown is SEM).

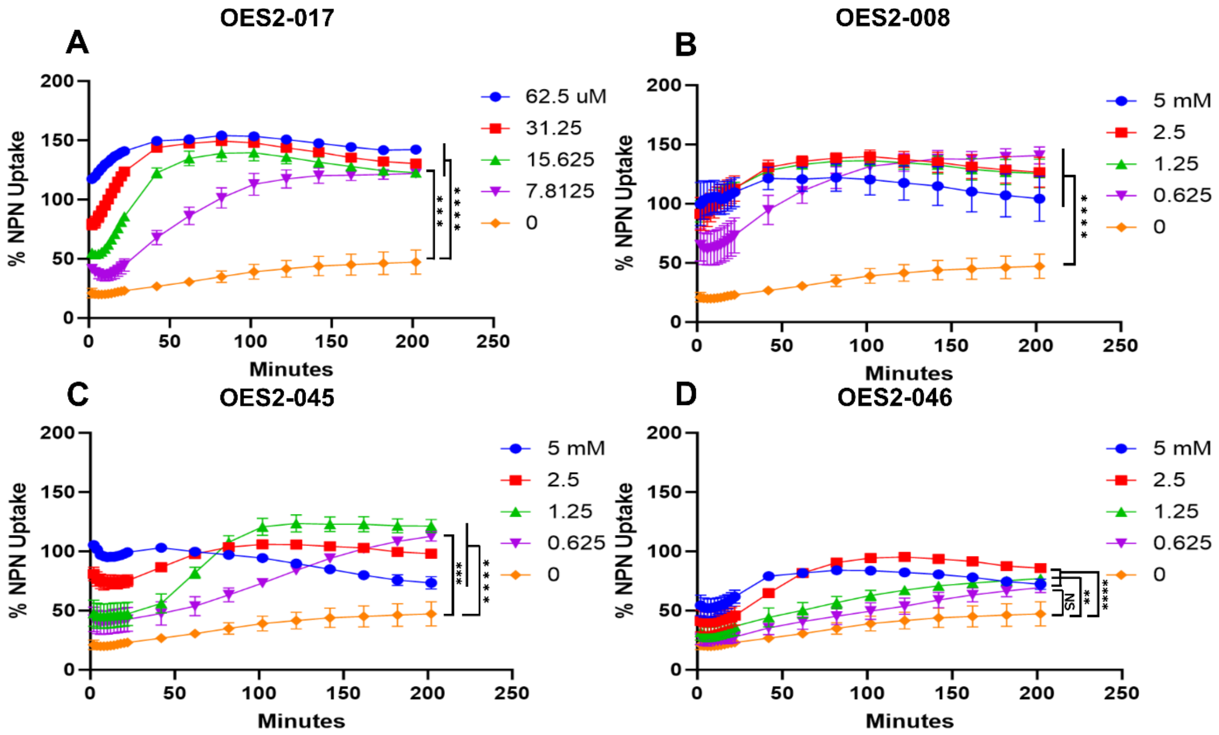


**Figure S10. Polyamine analogs increase the outer membrane permeability of *K. pneumoniae* to NPN.** A-D) A kinetic assay displaying % NPN uptake in *K. pneumoniae* (MKP103) induced by perturbation of the outer membrane by the polyamine analogs A) OES2-017, B) OES2-008, C) OES2-045, and D) OES2-046 (error bars shown as SEM). A one-way ANOVA analysis followed by a Brown-Forsythe and Welch test assessed the significance of the difference as indicated on the plot.

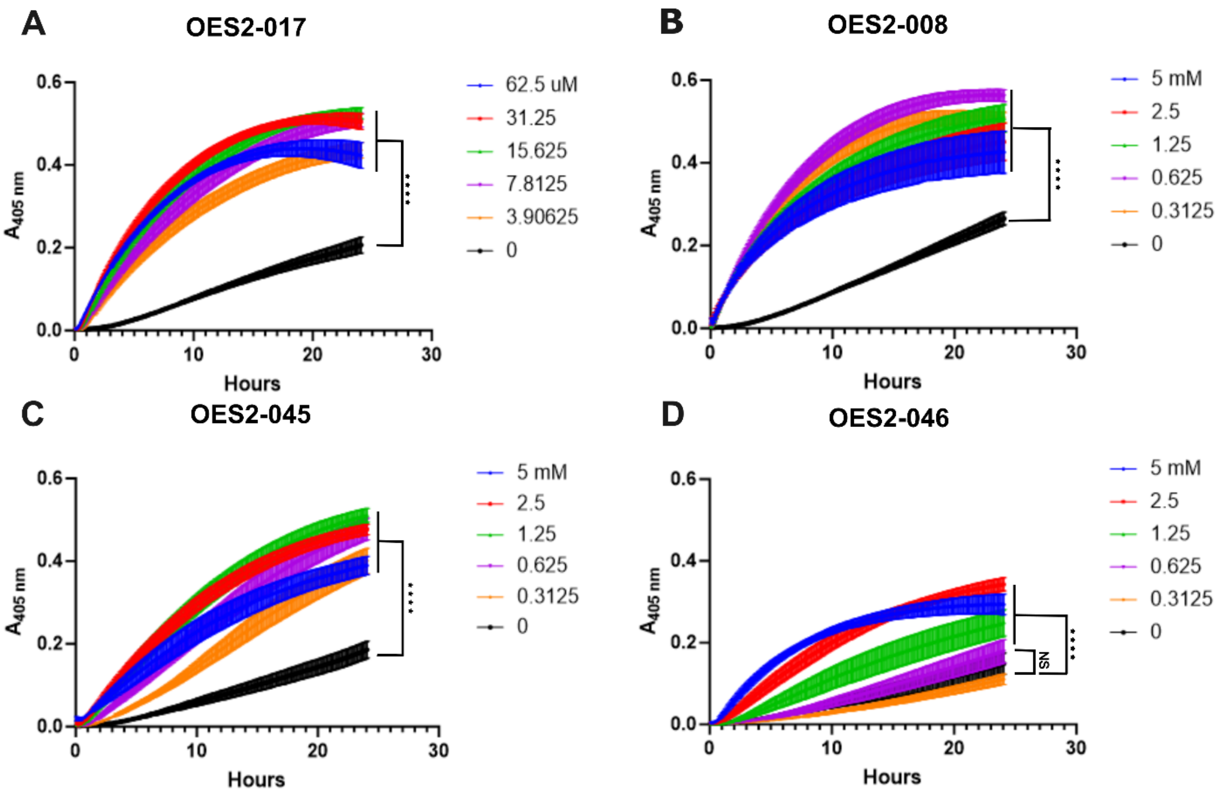


**Figure S11. Polyamine analogs increase the inner membrane permeability of *E. coli* ML-35 pBR322 to ONPG.** A-D) A kinetic assay displaying absorbance of ONPG hydrolysis in *E. coli* ML-35 pBR322 induced by perturbation of the inner membrane by the polyamine analogs A) OES2-017, B) OES2-008, C) OES2-045, D) and OES2-046.; error bars shown as SEM. A one-way ANOVA analysis followed by a Brown-Forsythe and Welch test assessed the significance of the difference as indicated on the plot.

**Table S1.** ***K. pneumoniae*** **diversity panel putrescine-azithromycin FICI values**

| Strain | FICI | Strain | FICI | Strain | FICI | Strain | FICI |
|---|---|---|---|---|---|---|---|
| MRSN 761403 | 1 | MRSN 583141 | 0.25 | MRSN 410359 | 0.375 | MRSN 21304 | 0.375 |
| MRSN 680172 | 0.625 | MRSN 583114 | 0.625 | MRSN 401050 | 0.625 | MRSN 20522 | 0.375 |
| MRSN 607210 | 0.75 | MRSN 581745 | 0.625 | MRSN 380979 | 0.5 | MRSN 19073 | 0.375 |
| MRSN 669510 | 0.515 | MRSN 613682 | 0.625 | MRSN 375436 | 0.375 | MRSN 18411 | 0.375 |
| MRSN 582610 | 1 | MRSN 752729 | 0.75 | MRSN 374613 | 0.375 | MRSN 16233 | 0.5 |
| MRSN 742743 | 0.375 | MRSN 564304 | 0.5 | MRSN 371351 | 0.375 | MRSN 16008 | 0.5 |
| MRSN 730567 | 0.25 | MRSN 562722 | 0.375 | MRSN 368320 | 0.375 | MRSN 15937 | 1 |
| MRSN 750877 | 0.375 | MRSN 560539 | 0.25 | MRSN 368001 | 1 | MRSN 15882 | 0.25 |
| MRSN 740795 | 0.25 | MRSN 546733 | 0.375 | MRSN 366562 | 0.375 | MRSN 15687 | 0.625 |
| MRSN 728987 | 0.375 | MRSN 539414 | 0.375 | MRSN 365679 | 0.625 | MRSN 15219 | 0.375 |
| MRSN 731029 | 0.375 | MRSN 526410 | 0.25 | MRSN 28893 | 0.5 | MRSN 14444 | 0.375 |
| MRSN 702325 | 0.25 | MRSN 518712 | 0.375 | MRSN 28887 | 0.375 | MRSN 13768 | 0.375 |
| MRSN 750999 | 0.375 | MRSN 517281 | 0.3125 | MRSN 28880 | 0.375 | MRSN 13761 | 0.375 |
| MRSN 750999 | 0.375 | MRSN 516635 | 0.375 | MRSN 28866 | 0.375 | MRSN 13748 | 0.375 |
| MRSN 736213 | 0.25 | MRSN 515432 | 0.375 | MRSN 28183 | 0.375 | MRSN 13726 | 0.75 |
| MRSN 681054 | 0.25 | MRSN 515247 | 0.3125 | MRSN 27989 | 0.5 | MRSN 7076 | 0.25 |
| MRSN 702261 | 0.375 | MRSN 513382 | 0.3125 | MRSN 27778 | 0.75 | MRSN 6778 | 0.375 |
| MRSN 699654 | 0.187 | MRSN 511348 | 0.375 | MRSN 27106 | 0.5 | MRSN 6031 | 0.375 |
| MRSN 672476 | 0.312 | MRSN 499958 | 0.375 | MRSN 25947 | 0.375 | MRSN 5881 | 0.75 |
| MRSN 676980 | 0.375 | MRSN 479404 | 0.375 | MRSN 25616 | 0.375 | MRSN 5741 | 0.625 |
| MRSN 669448 | 0.312 | MRSN 468268 | 0.375 | MRSN 25112 | 0.375 | MRSN 5613 | 0.375 |
| MRSN 614201 | 0.375 | MRSN 450199 | 0.375 | MRSN 25107 | 0.375 | MRSN 4815 | 0.375 |
| MRSN 599975 | 0.375 | MRSN 430414 | 0.5 | MRSN 22265 | 0.625 | MRSN 4759 | 0.375 |
| MRSN 572640 | 0.25 | MRSN 430405 | 0.75 | MRSN 22232 | 0.5 | MRSN 4111 | 0.25 |
| MRSN 591344 | 0.25 | MRSN 414780 | 0.375 | MRSN 21352 | 1 | MRSN 1912 | 0.75 |

**Table S2.** ***K. pneumoniae*** **Wild-Type and** ***waaD*****::Tn MIC comparison**

| **Compound** | **MIC wild-type** | **MIC *waaD*::Tn** | **Fold change** |
|---|---|---|---|
| Tetracycline | 32 | 16 | 2 |
| Chloramphenicol | 125 | ND* | ND |
| Gentamicin | 1 | 1 | 0 |
| Paromomycin | 512 | 512 | 0 |

*Not determined as the transposon contain a chloramphenicol resistance cassette
